# Stress-induced neuron remodeling reveals differential interplay between neurexin and environmental factors

**DOI:** 10.1101/462796

**Authors:** Michael P. Hart

**Affiliations:** Department of Genetics, University of Pennsylvania, Philadelphia PA 19104

## Abstract

Neurexins are neuronal adhesion molecules important for synapse maturation, function, and plasticity. Neurexins have been genetically associated with neurodevelopmental disorders including autism spectrum disorders (ASDs) and schizophrenia, but can have variable penetrance and phenotypic severity. Heritability studies indicate that a significant percentage of risk for ASD and schizophrenia includes environmental factors, highlighting the poorly understood interplay between genetic and environmental factors in pathogenesis of these disorders. The singular *C. elegans* ortholog of human neurexins, *nrx-1*, controls experience-dependent morphologic remodeling of a GABAergic neuron in adult males. Here I show that this GABAergic neuron’s morphology is altered in response to each of three environmental stressors (nutritional, heat, or genotoxic stress) applied during sexual maturation, but not during adulthood. Increased outgrowth of axon-like neurites following adolescent stress results from an altered morphologic plasticity that occurs upon entry into adulthood. Despite axonal remodeling being induced by each of the three stressors, only nutritional stress (starvation) impacts behavior and is dependent on neurexin*/nrx-1*. Heat or genotoxic stress during sexual maturation did not alter behavior despite inducing GABAergic neuron remodeling, and this remodeling was independent of neurexin/*nrx-1*. Remodeling induced by starvation stress was found be dependent on neuroligin/*nlg-1*, the canonical binding partner for neurexin/*nrx-1*, as well as the stress signaling transcription factors *FOXO*/*daf-16* and *HSF1*/*hsf-1*, each of which was also found to have unique roles in remodeling induced by heat and UV stress. The differential molecular mechanisms underlying GABAergic neuron remodeling in response to different stressors, and the disparate effects of stressors on behavior, is a novel paradigm for understanding how genetics, environmental exposures, and plasticity may contribute to brain dysfunction in ASDs and schizophrenia.

## INTRODUCTION

Autism spectrum disorders (ASDs) and schizophrenia are neurodevelopmental/neuropsychiatric disorders characterized by social and cognitive phenotypes that can be extremely heterogeneous in makeup and severity. Their complex etiology is a consequence of pathogenic roles of both genetic and environmental risk factors (Cattane, Richetto, & Cattaneo, 2018; Chaste & Leboyer, 2012; Hallmayer et al., 2011; Hertz-Picciotto, Schmidt, & Krakowiak, 2018; Sandin et al., 2014; Schmitt, Malchow, Hasan, & Falkai, 2014; Tick, Bolton, Happe, Rutter, & Rijsdijk, 2016). Exposure to stress is a specific environmental risk factor for ASDs and schizophrenia, and may influence the course of these disorders (Bishop-Fitzpatrick, Mazefsky, Minshew, & Eack, 2015; Fuld, 2018; Schmitt et al., 2014). Recent work on the genetics of ASDs and schizophrenia has implicated a number of genes involved in overlapping pathways to be associated with both disorders, including many neuronal/synaptic genes (Autism Spectrum Disorders Working Group of The Psychiatric Genomics, 2017; Waltereit, Banaschewski, Meyer-Lindenberg, & Poustka, 2014). However, the potentially important interplay between variants in genes associated with ASDs/schizophrenia and environmental exposures, including stress, is still not clearly defined at the cellular and molecular level.

Stress exposure impacts the brain in many ways that might explain the connection between stress and neurodevelopmental disorders. For example, changes in neuron morphology (especially of the dendrites and axons), which are known to be induced by stress, can mediate changes in neuron and circuit function by dictating connectivity potential. More than a decade of work has characterized the impact of stress on neuron morphology in the vertebrate brain, specifically with regard to atrophy and/or hypertrophy of excitatory dendrites in the hippocampus, amygdala, and cortex in response to stress (Cook & Wellman, 2004; McEwen, Nasca, & Gray, 2016; Vyas, Mitra, Shankaranarayana Rao, & Chattarji, 2002). Recent work has also identified stress-induced dendritic remodeling in GABAergic inhibitory interneurons in the same brain regions, in some cases mirroring the direction of dendrite remodeling of excitatory neurons (Gilabert-Juan, Bueno-Fernandez, Castillo-Gomez, & Nacher, 2017; Gilabert-Juan, Castillo-Gomez, Guirado, Molto, & Nacher, 2013; Gilabert-Juan, Castillo-Gomez, Perez-Rando, Molto, & Nacher, 2011).

However, much less is known about remodeling of neuronal axons in response to stres*s*, in large part due to the technical difficulty of performing such studies in the vertebrate brain. Stress has been found to alter the presynaptic dynamics of inhibitory neurons, indicating that axons can be impacted by stress (Czeh et al., 2018; McKlveen et al., 2016). Studies have tried to connect stress-induced changes in neuron morphology with changes in the functional output of neurons, namely relevant behavioral changes. While stress-induced dendritic hypertrophy of neurons in the rodent hippocampus have been shown to correlate with changes in spatial learning tasks, some studies have demonstrated that behavioral changes following stress can be separated from neuron remodeling events (Conrad, 2006; Conrad, LeDoux, Magarinos, & McEwen, 1999; Conrad, Ortiz, & Judd, 2017; Vyas et al., 2002). Thus, a direct connection between stress-induced dendritic remodeling and behavior has been difficult to fully define. The majority of vertebrate studies also lack the temporal resolution to analyze the dynamics of neuron morphologic changes following stress, and while the mechanisms controlling excitatory dendrite remodeling in response to stress have been studied (McEwen et al., 2016), those controlling remodeling in inhibitory neurons, or the interplay between excitatory and inhibitory remodeling, are still relatively unknown.

*Caenorhabditis elegans* has been utilized as a powerful and simple model for neuronal response to stress, including various environmental, internal/developmental, and aging stressors (Kagias, Nehammer, & Pocock, 2012). We recently described experience-dependent morphologic plasticity in the nematode *C. elegans*, where the GABAergic DVB neuron undergoes progressive experience-dependent outgrowth of axon-like neurites in adult males. This plasticity alters circuit connectivity to cause changes in male mating and defecation behaviors (Hart & Hobert, 2018). We identified the singular *C. elegans* ortholog of human neurexins, *nrx-1*, to be required in the DVB neuron for this neurite outgrowth. Neurexins are synaptic adhesion molecules with diverse functions in synaptic formation, maturation, maintenance, function, and plasticity (Sudhof, 2017), and have been genetically associated with both ASDs and schizophrenia (Gauthier et al., 2011; Kirov et al., 2009; Reichelt, Rodgers, & Clapcote, 2012). Therefore, neurexins are strong candidates for mediating changes in neuron morphology and behavior in response to environmental stress.

This *C. elegans* model of GABAergic morphologic plasticity provides a platform to investigate the impact of stress on GABAergic axonal morphology, including the temporal dynamics of remodeling. The results presented here demonstrate that stress, specifically during sexual maturation of the male nervous system (fourth larval stage, ‘adolescence’ (Snoek et al., 2014)), alters the morphology of the DVB GABAergic neuron, with outgrowth of axon-like neurites becoming more elaborate in early adulthood. Each of three distinct stressors – nutritional (4 or 18 hour fasting/starvation), heat (30 minutes at 37°C), or genotoxic (254 nm light set to 200 X100 μJ/cm^2^) – impact GABAergic neuron morphology, resulting in lasting morphologic changes. Despite this, each stressor differentially impacts downstream functional output and behavior with disparate dependence on *nrx-1*, its canonical binding partner neuroligin/*nlg-1* (also an ASD-associated gene), and two conserved stress responsive transcription factors (*FOXO*/*daf-16* and *HSF1*/*hsf-1)*. These findings demonstrate that different stressors alter axonal morphology and morphologic dynamics in GABAergic neurons with disparate effects on circuit dynamics and behavioral output, and provide an example of how underlying genetic defects (*nrx-1* or *nlg-1* mutation) can interact with different environmental stressors to affect distinct neuronal and behavioral responses. These results provide a platform for understanding the combinatorial impact of genetics and environment on risk for pathogenesis of neuropsychiatric disorders.

## RESULTS

### Adolescent stress alters GABAergic neuron morphology by altering morphologic plasticity in adulthood

The GABAergic DVB neuron is present in both male and hermaphrodite *C. elegans*, but undergoes experience-dependent morphological plasticity only in males (Hart & Hobert, 2018). In studying morphologic plasticity of the GABAergic DVB neuron in adult male *C. elegans* (**Fig.1A**), I noted that dietary history (fed vs. starved) appeared to affect DVB neurite outgrowth. To confirm this observation, I compared DVB neurite outgrowth between well-fed males and males cultivated without food for 18 hours beginning in the mid-L4 stage (**Fig.1B**). DVB in males ‘starved’ for 18 hours had increased total neurite length and neurite junctions (a proxy for the number of neurite branches calculated by Simple Neurite Tracer plugin for ImageJ/FIJI) compared to fed males (**Fig.1C-E**.). To determine if this effect was starvation-specific or a more general neuronal response to stress, fed males were subjected to heat or genotoxic stress (ultraviolet light (UV) exposure) at mid-L4 (**Fig.1B**). 18 hours after stress induction, both heat- and UV-stressed males had increases in DVB neurite outgrowth compared to unstressed animals (**Fig.1C-E**). Thus, three different stressors applied during the mid-L4 stage impacted DVB neuron morphology in a similar manner.

**Figure 1.**
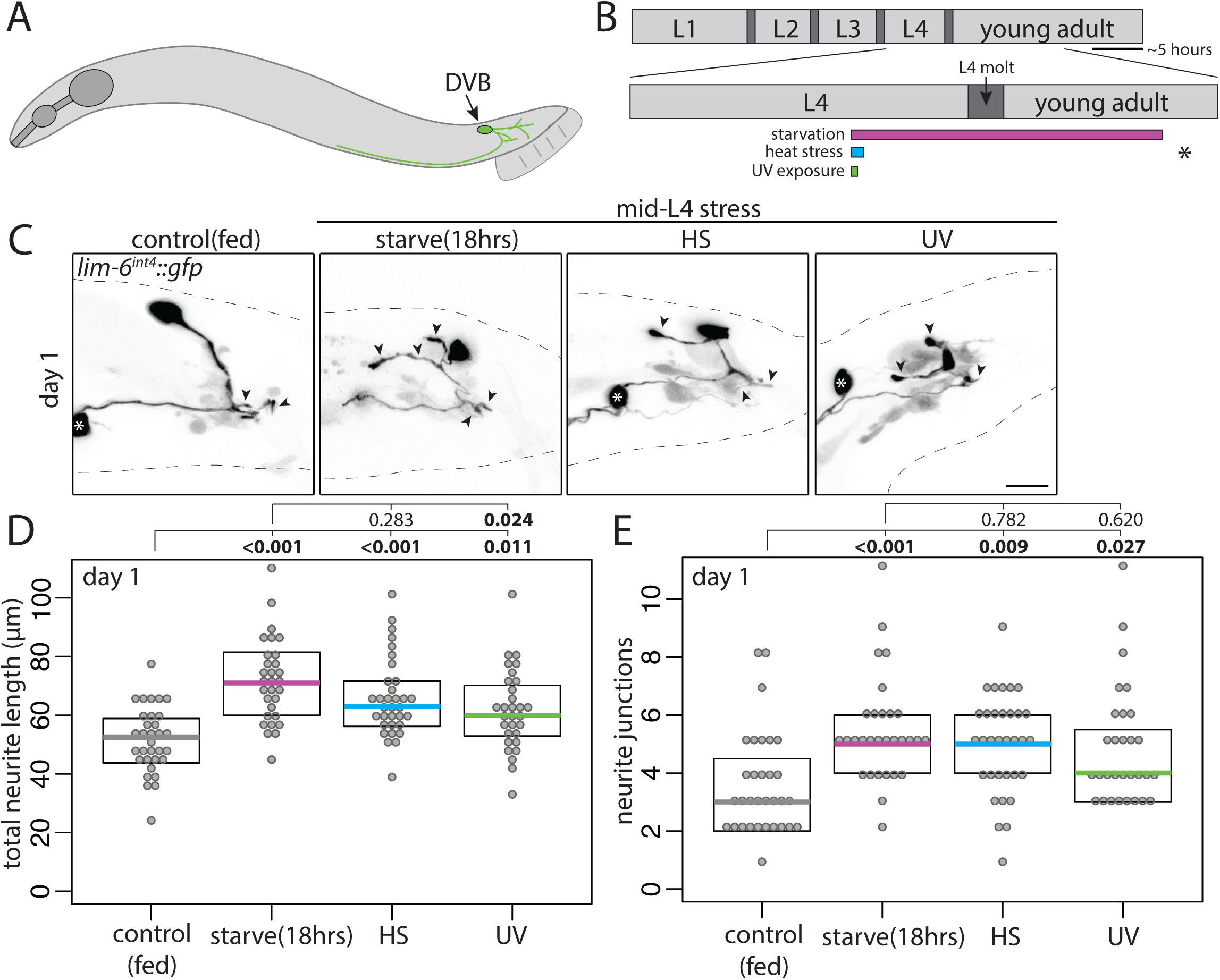
Multiple stressors during adolescence induce neurite outgrowth of the DVB neuron in adult male *C. elegans*. **A)** Illustration of male *C. elegans* with DVB neuron and neurites depicted. **B)** Timeline of post-embryonic development of male *C. elegans* with inset of L4 to adult period. Boxes below timeline indicate timing of different stressors labeled for each panel; grey – control/fed, magenta – starvation/fasting, teal – heat stress, green – UV exposure. Asterisk indicates time of confocal imaging and DVB neurite quantification. **C)** Confocal z-stack maximum projections of DVB neuron in control and mid-L4 stressed males at day 1 of adulthood (arrowheads indicate DVB neurites, white asterisk indicates soma of PVT neuron, scale bar = 10μm). Quantification of (**D**) total neurite length and (**E**) neurite junctions for DVB in control and mid-L4 stressed males at day 1 (Each dot represents one worm; colored bar, median; boxes, quartiles. Comparison using one-way ANOVA and post-hoc Tukey HSD, *P* values shown above plots with bold showing significance (*P*<0.05). True for all subsequent Figures).

The L4 stage, which is analogous to vertebrate adolescence, is when sexual maturation of the nervous system and other sexual tissues occurs, followed by the L4 molt, a period of quiescence prior to adulthood. Previous work demonstrates that transient fasting of a few hours has similar impact to 18-20 hours of starvation on male muscle and neuron excitability (LeBoeuf, Guo, & Garcia, 2011). To test the impact of timing during L4/L4 mold and length of starvation stress on DVB neuron morphology in adolescence, males were stressed early in the L4 stage (**Fig.2A**). Early-L4 heat stress and 4 hour fast both resulted in increases in DVB neurite outgrowth in day 1 males (**Fig.2B,C,D(last panel))**. UV exposure at early L4 resulted in the majority of males arresting before or during the L4 molt, precluding analysis of this stressor at the early L4 timepoint.

**Figure 2.**
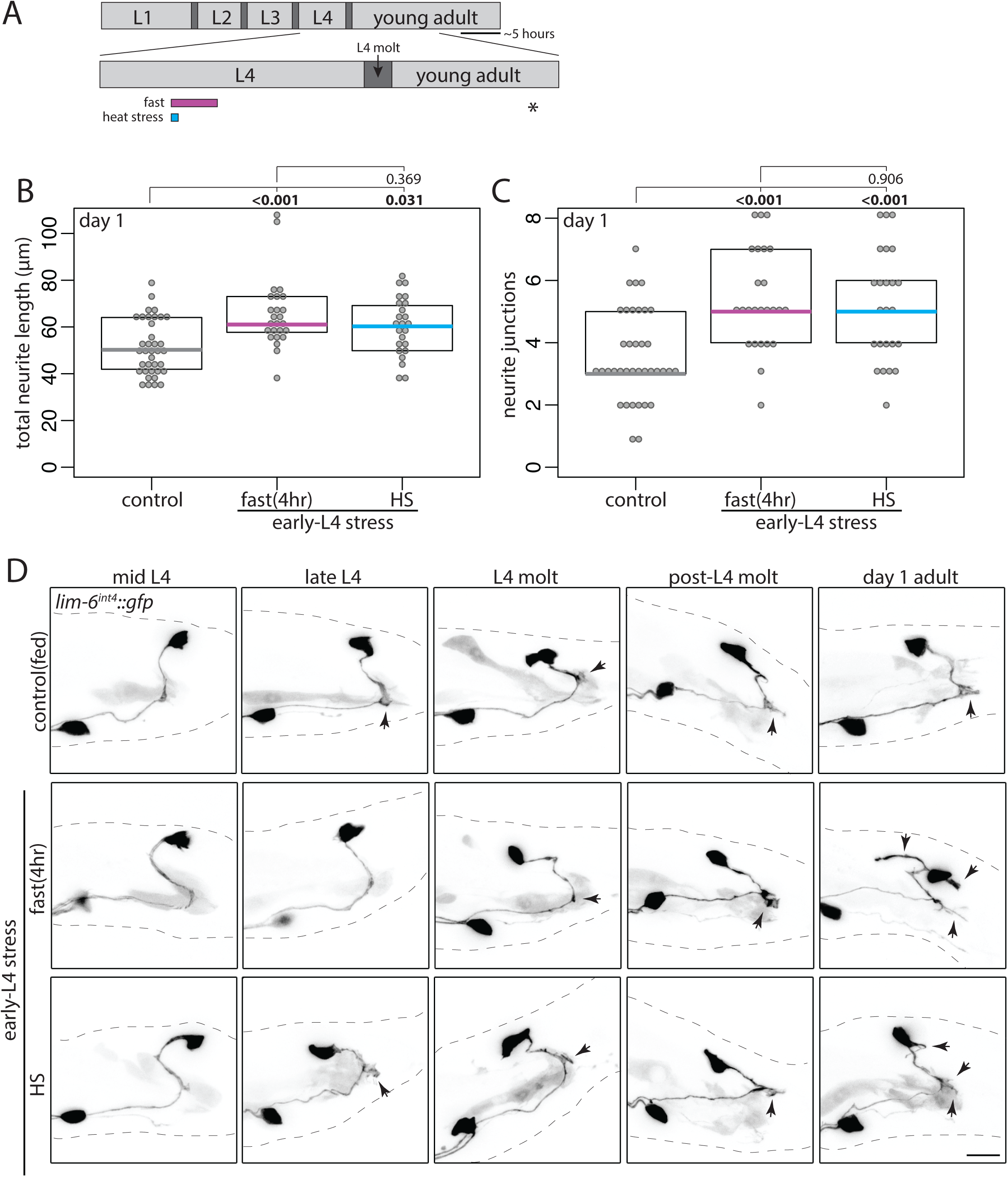
Adolescent stress alters DVB neurite outgrowth dynamics in adulthood. **A)** Diagram of L4 to adult period with boxes depicting stressors applied in early L4. Quantification of (**B**) total neurite length and (**C**) neurite junctions for DVB in control and early-L4 stressed males at day 1 of adulthood corresponding confocal maximum projections are last panels of (**D**). **D)** Confocal z-stack maximum projections of DVB neuron in control and early-L4 stressed males at different times throughout the L4 stage into early adulthood (10 males observed for each timepoint and condition)(scale bar = 10μm). Arrowheads indicate single neurites before, during, and after L4-molt.

Two explanations could account for these stress-induced morphologic changes in DVB neurite outgrowth. One possibility is that neurite outgrowth begins earlier in response to stress, such that by the first day of adulthood males have more DVB neurites. Alternatively, DVB neurite outgrowth may start at the same time as in unstressed males, but occur with different dynamics, resulting in adult males with increased neurites. To distinguish between these possibilities, I monitored DVB morphology in unstressed and in early-L4 stressed males at different timepoints from mid L4 into day 1 of adulthood (**Fig.2A,D**). Strikingly, despite stress being applied during early L4, DVB morphology was not different between non-stressed and stressed males until after the L4 molt and entry into adulthood (**Fig.2D**). At late-L4 and L4 molt, males had either no neurites or a single short neurite, regardless of stress. Only after L4 molt did differences begin to appear (**Fig.2D**), with stressed males having increased DVB neurites. This result supports the second explanation, that adolescent stress does not change the timing of DVB neurite outgrowth, but rather changes the degree of neurite outgrowth occurring in adult males. These results suggest that adolescent stress exposure changes the dynamics of DVB morphologic plasticity in adulthood, and that even a short fasting period of 4 hours during adolescence can alter adult GABAergic neuron morphology.

To test if stress-induced alterations to DVB neuron morphology were specific to the adolescent stage or if stress in early adulthood could have the same impact, male worms were exposed to stress shortly following entry into adulthood (**Fig.3A**). Surprisingly, the stress in early adulthood did not increase DVB neurite outgrowth (**Fig.3A-C**). In fact, 18 hour starvation in adult males resulted in a small, but significant, decrease in DVB total neurite length (**Fig.3A-C**). These results suggest that the increase in DVB neurite outgrowth in response to stress is specific to the L4 adolescent stage. Are stress-induced DVB morphologic changes long-lasting or a temporary and reversible morphologic stress response? DVB morphology was analyzed at day 3 of adulthood following mid-L4 stress (**Fig.3A**). Day 3 of adulthood corresponds to the end of peak male *C. elegans* reproductive capability (Guo, Navetta, Gualberto, & Garcia, 2012). Remarkably, all 3 stressors resulted in increases in at least one parameter of DVB neurite outgrowth more than 60 hours after stress (**Fig.3D,E**). In UV-exposed males both neurite length and junctions were increased, while in heat stressed males only neurite length was increased, and in starved males only neurite junctions were increased (**Fig.3D,E**). These results demonstrate that although all stressors increase neurite outgrowth at day 1, different stresses may impact DVB morphology in slightly different ways in the long term (**Fig.3D,E**). Thus, adult DVB neuron morphology is altered by adolescent stress as a consequence of enhanced morphologic plasticity in adulthood, producing long-lasting changes to GABAergic neuron morphology.

**Figure 3.**
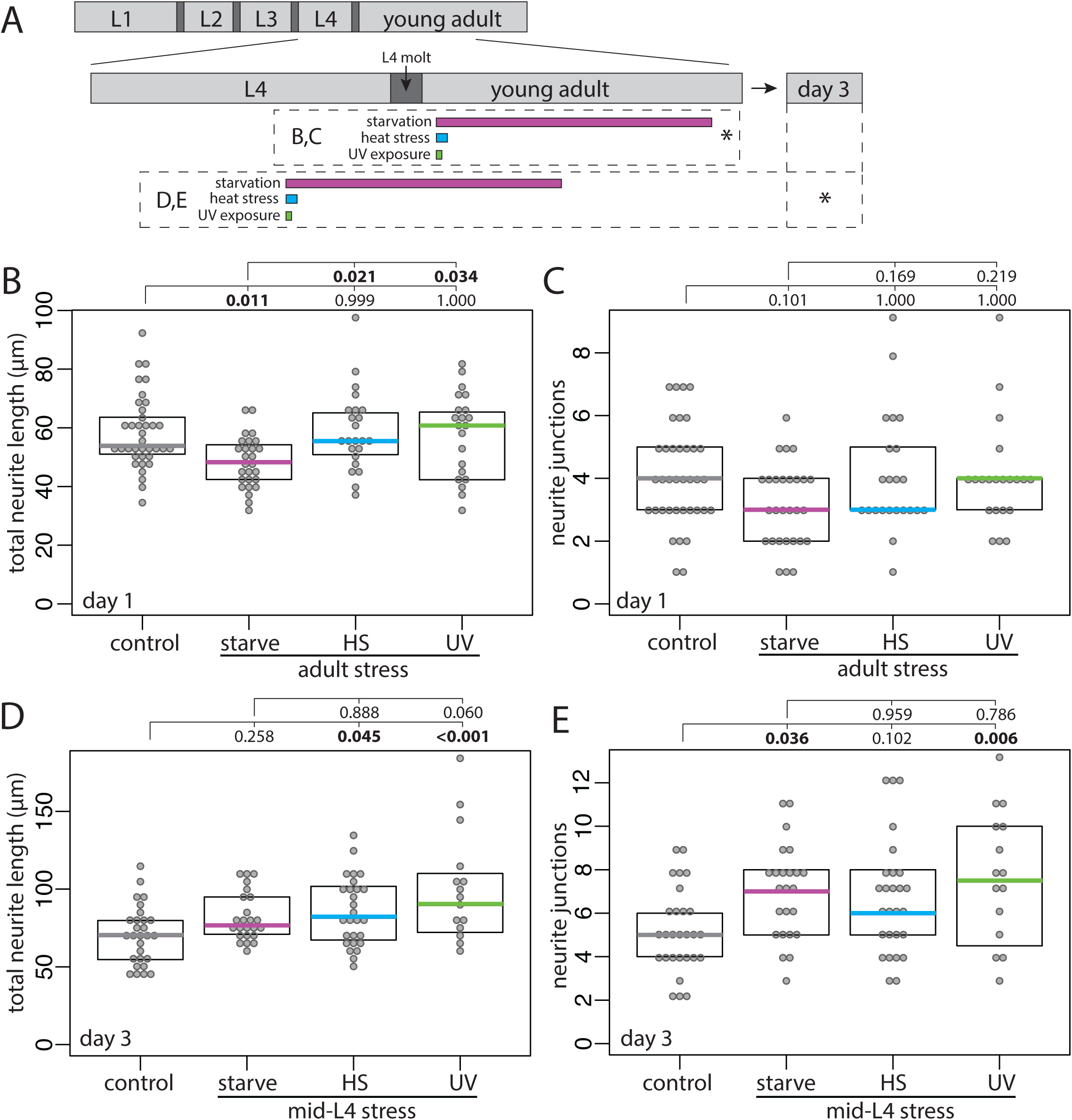
Early adulthood stress does not alter DVB morphoplogy, but adolescent stress results in sustained changes to DVB morphology into adulthood. **A)** Diagram of L4 to adult period, including inset indicating day 3 of adulthood. Boxes below timeline indicate timing of stressors, asterisk indicates time of confocal imaging and quantification. Quantification of (**B**) total neurite length and (**C**) neurite junctions for DVB in control and post-L4 molt stressed males at day 1. Quantification of (**D**) total neurite length and (**E**) neurite junctions for DVB in control and mid-L4 stressed males at day 3.

### Adolescent stressors differentially impact DVB function in male spicule protraction behavior and DVB presynaptic number and morphology

The functional impact of DVB morphologic plasticity in adulthood was previously characterized on a number of behaviors, including male spicule protraction behavior (Hart & Hobert, 2018). The spicules (**Fig.4A**) protract out of the male tail to penetrate the hermaphrodite vulva during copulation, and are therefore required for successful sperm transfer (Garcia, Mehta, & Sternberg, 2001). Males exposed to aldicarb, an acetylcholinesterase inhibitor, protract their spicules due to acetylcholine (ACh) buildup at the neuromuscular synapses between the cholinergic SPC neuron and the spicule protraction muscles (**Fig.4B**)(Garcia et al., 2001). The GABAergic DVB neuron synapses onto the SPC neuron as well as onto the spicule protractor muscles to inhibit spicule protraction (**Fig.4B**). Therefore, DVB neurite elaboration, which adds synapses specifically onto these targets (Hart & Hobert, 2018), results in increased SPC and spicule protractor inhibition, and a delay in the time to aldicarb-induced spicule protraction. This increased time to spicule protraction serves as a readout for the amount of DVB GABAergic input (Hart & Hobert, 2018).

**Figure 4.**
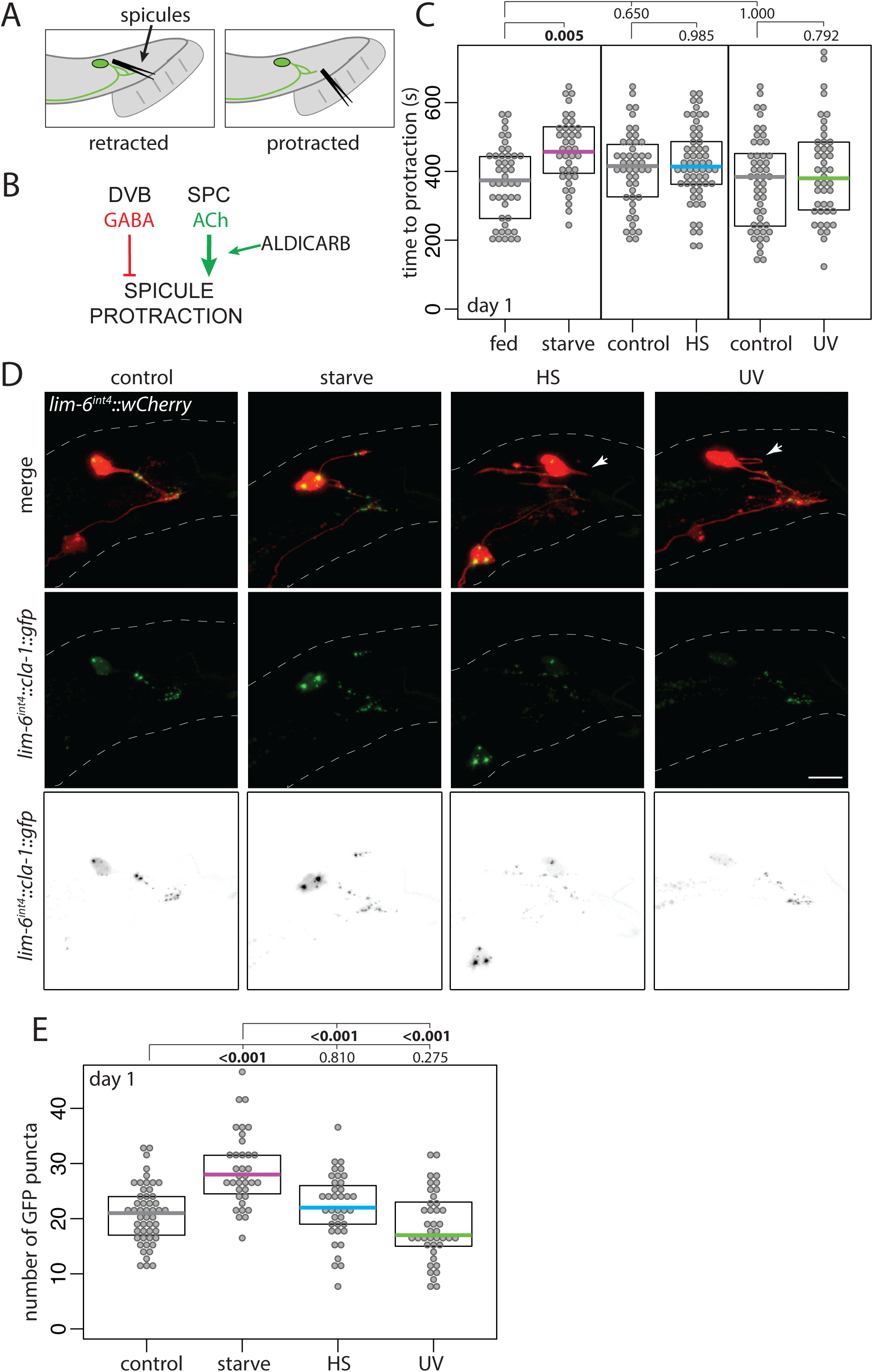
Adolescent stress differentially impacts adult spicule protraction behavior and number of DVB presynaptic active zone sites. **A)** Cartoon of *C. elegans* male tail with DVB and spicules depicted in retracted and protracted positions. **B)** Logic diagram of the male *C. elegans* spicule protraction circuit involving DVB GABAergic and SPC cholinergic signaling, showing location of action of the acetylcholinesterase inhibitor aldicarb. **C)** Time to protraction of spicules in control and mid-L4 stressed males on aldicarb plates at day 1 of adulthood. **D)** Confocal z-stack maximum projections of DVB neuron and DVB presynaptic puncta (*lim-6^int4^::cla-1::gfp*) in control and mid-L4 stressed males at day 1 of adulthood (arrowheads indicate DVB neurites with no GFP signal, scale bar = 10μm). (**E**) Number of GFP puncta in DVB neurites in control and mid-L4 stressed males at day 1.

To test if the outgrowth of DVB neurites induced by adolescent stress impacts the function and plasticity of DVB and the downstream spicule protraction behavior, unstressed and mid-L4-stressed males were exposed to aldicarb at day 1 of adulthood immediately following starvation stress (**Fig.4C**). Starved males had increased time to spicule protraction on aldicarb compared to fed males (**Fig.4C**), suggesting that the elaborated DVB neurites result in functionally enhanced GABAergic tone. In contrast, males exposed to either heat stress or UV had no change in time to spicule protraction in comparison to unstressed males (**Fig.4C**). These results suggest that although all of the stressors result in increased outgrowth of DVB neurites, the functional impact of these neurites is different between stressors – it is only starvation-induced DVB neurites that have an effect on spicule protraction behavior.

The above data demonstrate that elaborated DVB neurites induced by stress may not have the same impact on circuit connectivity, potentially explaining the disparate behavioral phenotypes. To explore this possibility, I analyzed DVB pre-synaptic puncta in unstressed and mid-L4 stressed males. GFP-tagged Clarinet1 (CLA-1), a piccolo/bassoon-like protein involved in synaptic vesicle release localizes to active zones (Kurshan et al., 2018; Xuan et al., 2017), while GFP-tagged RAB-3, a pre-synaptic GTPase that marks vesicle clusters, shows overlapping localization in the synaptic bouton (Kurshan et al., 2018). DVB-specific expression of CLA-1::GFP localized to bright distinct puncta on DVB processes, allowing visualization and quantification of pre-synaptic sites on neurites by counting puncta (**Fig.4D**). Day 1 males exposed to mid-L4 starvation stress had more CLA-1::GFP puncta on DVB neurites compared with non-stressed controls and heat or UV stressed males (**Fig.4D,E**). Localization of puncta within neurites did not appear different between conditions, except with some heat and UV-stressed males having neurites without any discernible GFP signal, which never occurred in non-stressed or starvation-stressed males (**Fig.4D**). GFP::RAB-3 expression in DVB was more diffuse with fewer distinct puncta that could not be quantified by counting. In day 1 starved males, GFP-tagged RAB-3 localized to bright puncta on all DVB neurites, compared with slightly dimmer and more diffuse localization in day 1 non-stressed males, perhaps indicating more mature synaptic structures following starvation that may correspond to the additional CLA-1::GFP puncta observed (**Fig.5A**). DVB presynaptic puncta in heat stressed and UV exposed males appeared dimmer and more diffuse, again with some neurites appearing very dim/diffuse or with no GFP signal (arrowheads, **Fig.5A**). To quantify the potential differences, ‘puncta’ were outlined through application of consistent thresholding on confocal z-stack maximum projections using the particle analysis feature in FIJI. A region of interest was drawn around DVB neurites, and particles (puncta) analyzed for various measurements. While there was no difference in the number of puncta between unstressed and any of the stressed males (data not shown), the average area of puncta in starved males was significantly increased compared to controls, while heat stressed and UV exposed males showed no difference from controls (**Fig.5B**). Additionally, the mean intensity of puncta was also increased in starved males compared to controls, but not males exposed to heat or UV (**Fig.5C**). Taking the results of both pre-synaptic markers, starved males have more presynaptic active zones with larger and brighter DVB vesicle clusters than controls, a difference not observed in males exposed to heat or UV stress, which may begin to explain the functional differences observed in behavioral output between the stressors.

**Figure 5.**
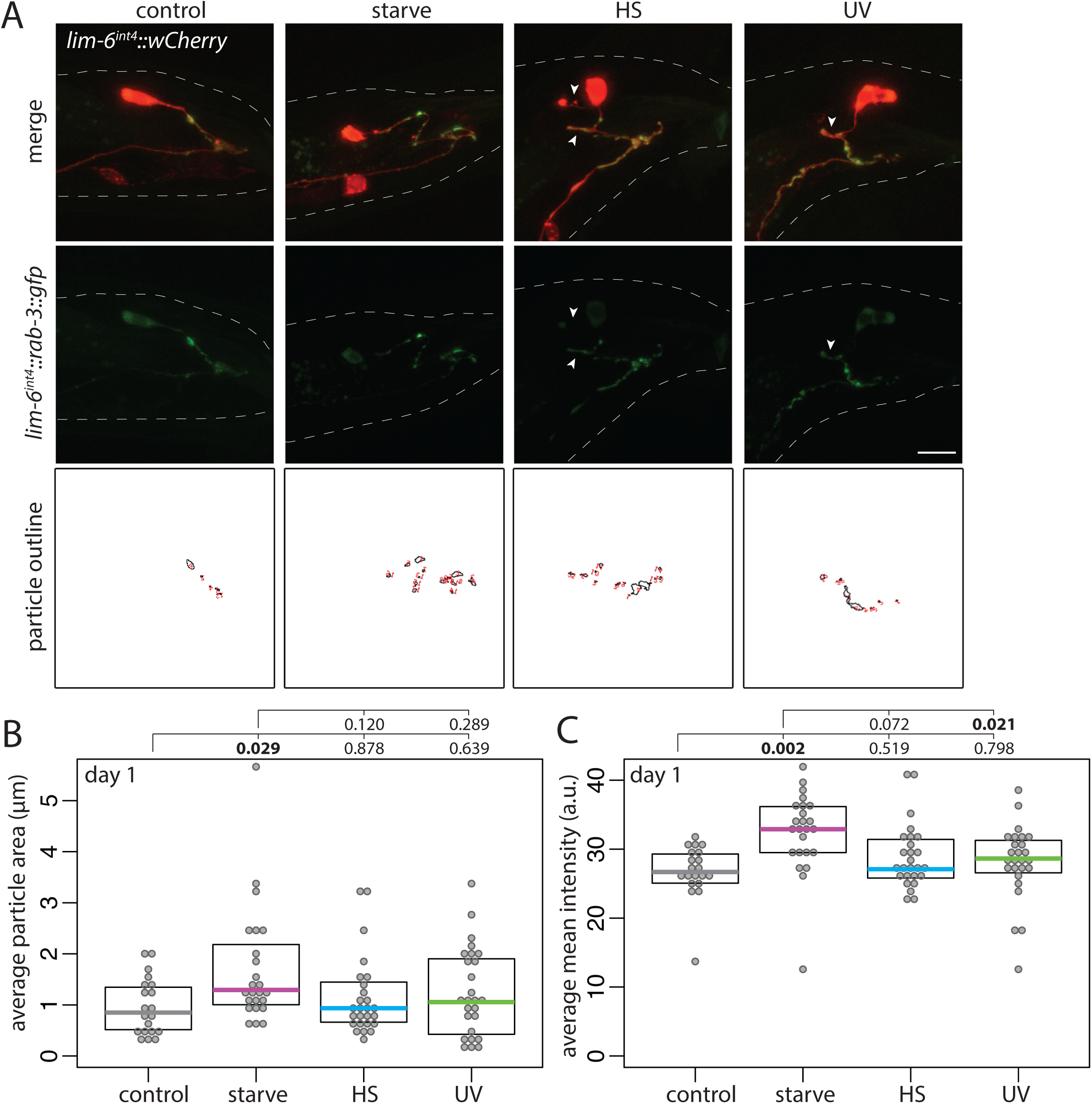
Adolescent stress differentially impacts DVB presynaptic vesicle cluster morphology. **A)** Confocal z-stack maximum projections of DVB neuron and DVB presynaptic puncta (*lim-6^int4^::gfp::rab-3*) in fed control and mid-L4 stressed males at day 1 of adulthood (arrowheads indicate DVB neurites with little/no gfp or diffuse gfp, scale bar = 10μm). Identically thresholded z-stack maximum projections of the *gfp::rab-3* channel created particle outlines of synaptic puncta on DVB neurites for analysis of intensity and other measures. Quantification of *lim-6^int4^::gfp::rab-3* (**B**) average particle area and (**C**) average mean intensity in control and mid-L4 stressed males at day 1.

### The *C. elegans* neurexin ortholog, *nrx-1*, is required for starvation-induced DVB neurite outgrowth

DVB neurite outgrowth in early adulthood is dependent on the ortholog of the synaptic adhesion molecule neurexin, *nrx-1* (Hart & Hobert, 2018). To determine if mid-L4 stress-induced DVB neurite outgrowth is also dependent on *nrx-1*, I analyzed neurite outgrowth in *nrx-1* mutant males. Two *nrx-1* alleles that strongly reduce gene function but differ in their effects on the two major *nrx-1* isoforms were used: a large deletion that removes the entire short isoform and the transmembrane and intracellular portion of the long isoform (which was recently described to have phenotypes indistinguishable from those of worms with a deletion covering the entire *nrx-1* locus (Kurshan et al., 2018), or a point mutation resulting in a premature stop codon affecting only the long isoform (**Fig.6A**). As previously observed, loss of *nrx-1* did not impact DVB neurite outgrowth at day 1 under fed conditions (**Fig.6B-G**)(Hart & Hobert, 2018). However, the increase in DVB neurite outgrowth observed in wild-type males following mid-L4 starvation did not occur in males with either *nrx-1* loss of function allele, with DVB neurites in starved *nrx-1* mutant males similar to fed wild-type males (**Fig.6B-G**). Remarkably, following either heat stress or UV exposure, animals harbouring either *nrx-1* loss of function allele showed increases in DVB neurite outgrowth compared to wild-type non-stressed males, similar to levels of outgrowth seen in heat-stressed or UV-exposed wild-type animals (**Fig.6B-G**). These results imply that while DVB neurite outgrowth and morphologic plasticity in response to starvation is dependent on *nrx-1*, heat and UV-induced DVB neurite outgrowth is not. Despite all three stressors resulting in increased DVB neurite outgrowth, a role for *nrx-1* is uniquely required for DVB response to starvation.

**Figure 6.**
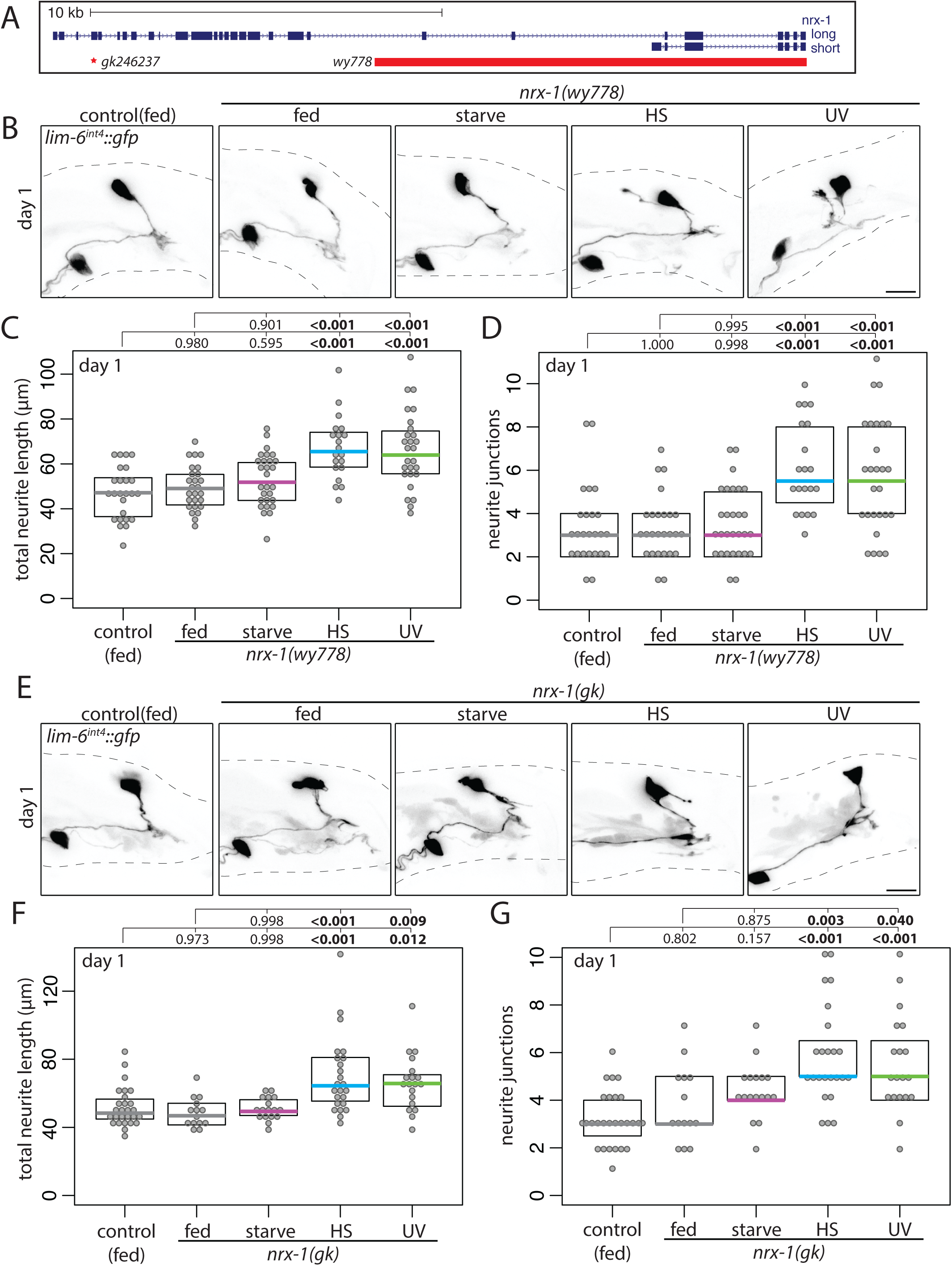
Neurexin is required for DVB neurite outgrowth following adolescent starvation stress. **A)** Genomic locus of *nrx-1* shown 5’ to 3’ with long and short isoforms included, red bar indicates region deleted in *nrx-1(wy778)* and red asterisk indicates point mutation of early stop in *nrx-1(gk246237)* referred to in Figures as *nrx-1(gk)*. **B)** Confocal z-stack maximum projections of DVB neuron in fed control and fed and mid-L4 stressed *nrx-1(wy778)* males at day 1 of adulthood (scale bar = 10μm). Quantification of (**C**) total neurite length and (**D**) neurite junctions for DVB in fed control and fed and mid-L4 stressed *nrx-1(wy778)* males at day 1. **E)** Confocal z-stack maximum projections of DVB neuron in fed control and fed and mid-L4 stressed *nrx-1(gk246237)* males at day 1 of adulthood (scale bar = 10μm). Quantification of (**F**) total neurite length and (**G**) neurite junctions for DVB in fed control and fed and mid-L4 stressed *nrx-1(gk246237)* males at day 1.

To test if *nrx-1* also has an impact on the functional output of DVB neurite outgrowth following adolescent stress, males with *nrx-1* deletion were exposed to mid-L4 stress and the aldicarb spicule protraction assay. Non-stressed *nrx-1* mutants were not different than non-stressed wild-type animals in time to protraction on aldicarb at day 1 (**Fig.7**). Whereas starved wild-type animals were slower to protract spicules on aldicarb than fed wild-type animals, no difference in time to protraction on aldicarb between fed and starved *nrx-1* mutants was observed (**Fig.7**), implicating *nrx-1* in both the underlying mechanism responsible for DVB neurite outgrowth and behavioral response to starvation. *nrx-1* males exposed to UV stress had no difference in the time to protraction compared to non-stressed *nrx-1* males (**Fig.7**), similar to UV stress not impacting time to protraction in wild-type animals. Unexpectedly, *nrx-1* males exposed to heat stress had an increase in the time to protraction on aldicarb compared to non-stressed *nrx-1* males (**Fig.7**), a difference not observed in wild-type males exposed to heat stress (**Fig.4C**). This suggests that even though *nrx-1* plays no role in heat stress-induced DVB morphologic changes, it may play a role in modifying the functional output of DVB or the spicule protraction circuit in response to heat stress, although how and where this occurs will require further investigation. Together, these results show that both the morphologic and functional plasticity of DVB in response to starvation are dependent on *nrx-1*, and demonstrate that different types of stress can have unique behavioral outcomes and depend on different underlying mechanisms.

**Figure 7.**
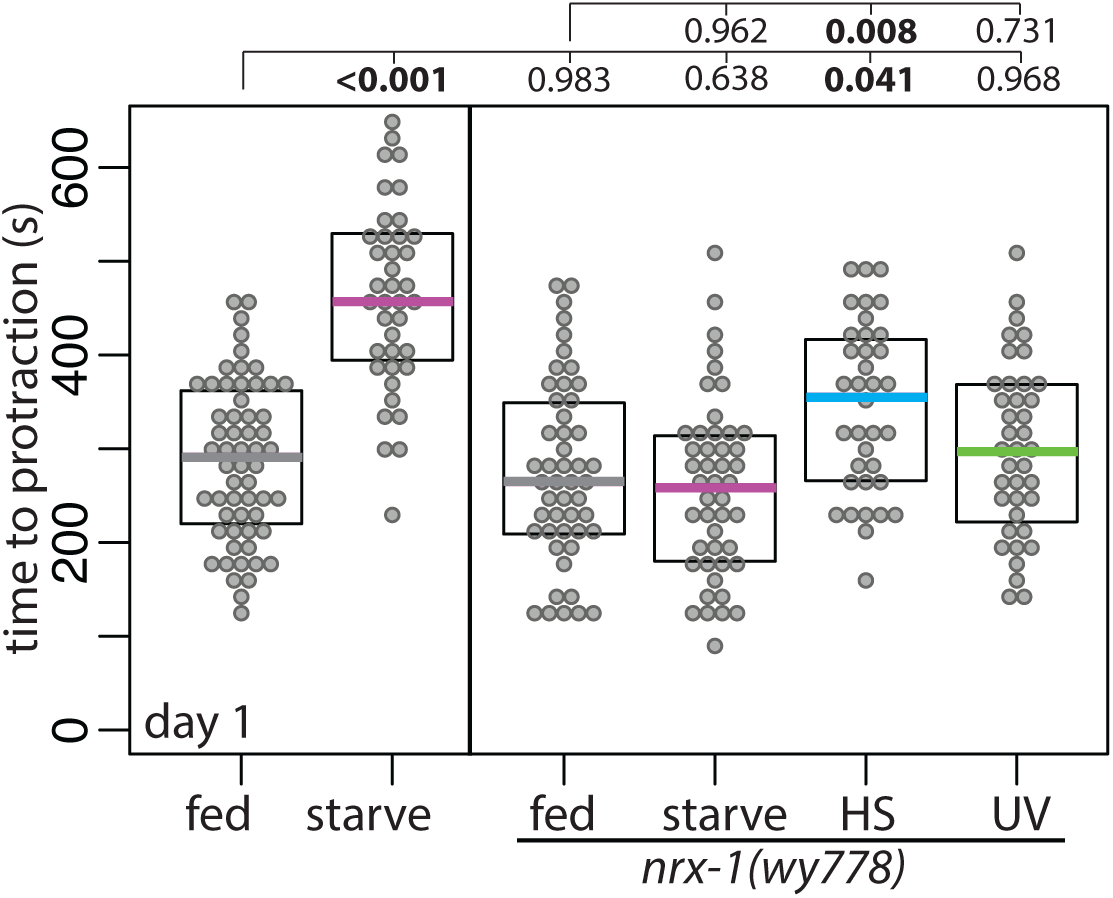
*nrx-1* is required for adolescent starvation-induced changes in adult spicule protraction behavior. Time to protraction of spicules in fed control, starved control (repeated data from Fig.3C, for comparison), and mid-L4 control and stressed *nrx-1(wy778)* males on aldicarb plates at day 1 of adulthood.

Neurexin was previously found to control experience-dependent DVB neurite outgrowth in adult males in a cell autonomous manner, with DVB-specific NRX-1 expression rescuing DVB neurite outgrowth and spicule protraction phenotypes of *nrx-1* mutant males at day 3 of adulthood (Hart & Hobert, 2018). Surprisingly, NRX-1 expression in DVB was unable to rescue the neurite outgrowth defect in *nrx-1* mutants following mid-L4 starvation (also observed for a GFP tagged NRX-1, data not shown)(**Fig.8A-C**). NRX-1 is expressed in all neurons and some muscles in *C. elegans* and recent work has described cell non-autonomous functions of NRX-1 (Philbrook et al., 2018; Tong et al., 2017). NRX-1 expression in all neurons using a pan-neuronal promoter (*ric-19p*(Stefanakis, Carrera, & Hobert, 2015)), did not significantly alter DVB neurite outgrowth in non-stressed males, but rescued the loss of DVB neurite outgrowth in *nrx-1* mutants following mid-L4 starvation (**Fig.8A-C**). Thus, NRX-1 expression in DVB alone cannot rescue deficits in starvation-induced neurite outgrowth, and NRX-1 expression in additional neurons is required for induction of neurite outgrowth following starvation. It is likely that NRX-1 is required in DVB to promote neurite outgrowth in a cell autonomous manner given its role in experience-dependent neurite outgrowth (Hart & Hobert, 2018), but that it is also needed in additional neurons to rescue deficits in the starvation-induced phenotype. It is possible that NRX-1 is required in neurons upstream of DVB to signal starvation stress, or that it is required for normal synaptic transmission at upstream and DVB sites that are required for activity-dependent DVB neurite outgrowth, or a combination of these two possibilities. Further work will be required to identify the specific neurons in which NRX-1 expression is required for stress signaling and subsequent starvation-induced DVB neurite outgrowth.

**Figure 8.**
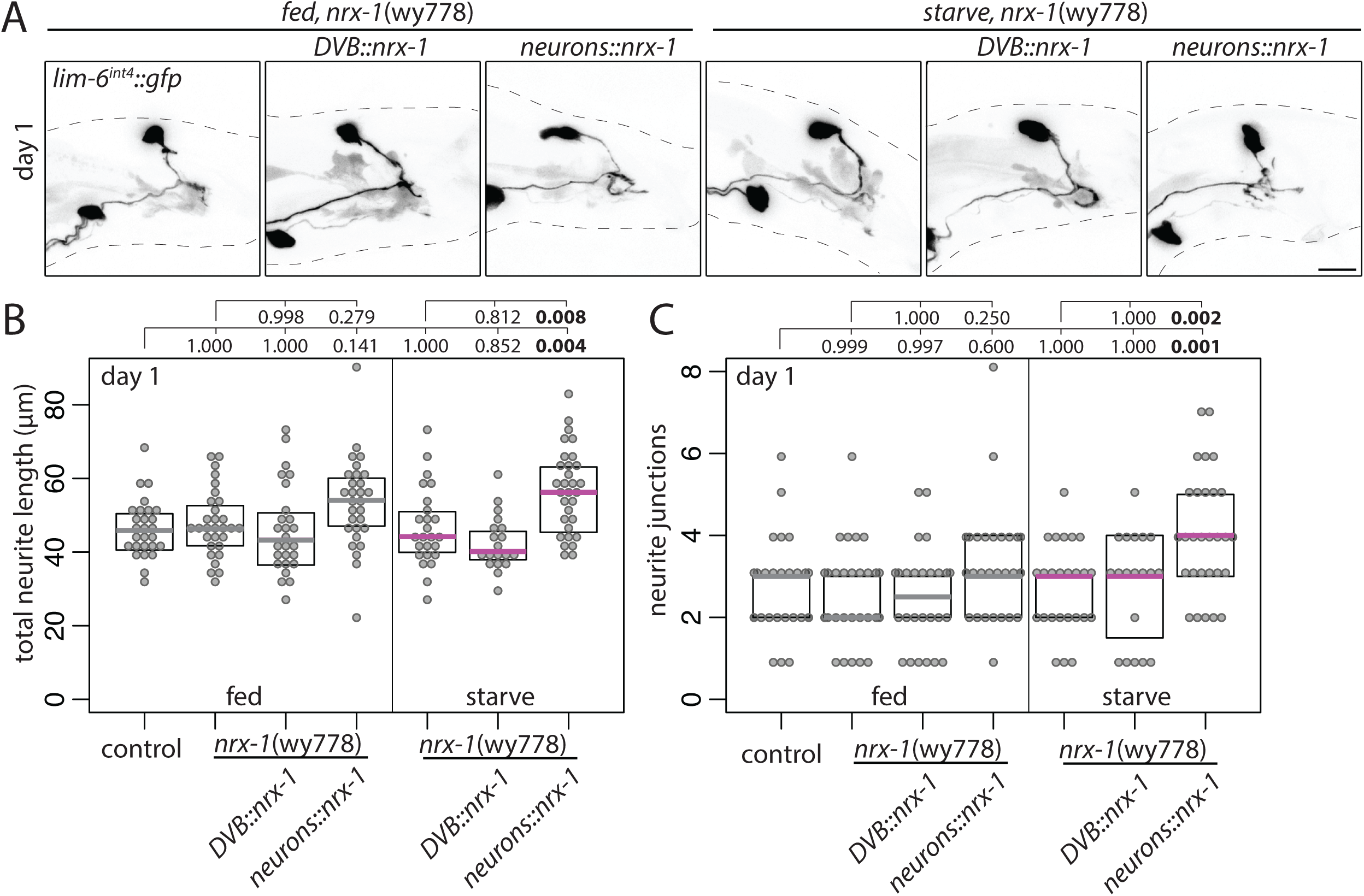
Neurexin expression in all neurons, but not in DVB alone, rescues loss of starvation-induced DVB neurites in *nrx-1* mutants. A) Confocal z-stack maximum projections of DVB neuron in fed and mid-L4 starved *nrx-1(wy778)* males with control, *DVB::nrx-1* (*lim-6^int4^::birA::nrx-1*), or *neurons::nrx-1* (*ric-19^prom6^::birA::nrx-1*) transgenes at day 1 of adulthood (scale bar = 10μm). Quantification of (**B**) total neurite length and (**C**) neurite junctions for DVB in fed control and fed and mid-L4 starved *nrx-1(wy778)* males with control, *DVB::nrx-1* (*lim-6^int4^::birA::nrx-1*), or *neurons::nrx-1* (*ric-19^prom6^::birA::nrx-1*) transgenes males at day 1 of adulthood.

### Neuroligin/*nlg-1* promotes starvation-induced DVB neurite outgrowth together with *nrx-1*

Neuroligins are synaptic adhesion molecules that interact with neurexins at synapses, commonly serving as the post-synaptic ligands for neurexins (Maro et al., 2015; Tong, Hu, Liu, Anderson, & Kaplan, 2015; Tu, Pinan-Lucarre, Ji, Jospin, & Bessereau, 2015). DVB experience-dependent neurite outgrowth in adult males requires NRX-1 in DVB to promote neurite outgrowth, while the *C. elegans* singular ortholog of neuroligins, NLG-1, restricts DVB neurite outgrowth from post-synaptic targets (Hart & Hobert, 2018). Does NLG-1 play a role in adolescent stress-induced DVB neurite outgrowth? DVB neurite outgrowth was not increased in day 1 *nlg-1* mutant males following starvation or heat stress, but did increase after UV exposure (**Fig.9A-C**), suggesting that NLG-1 is required for DVB neurite outgrowth induced by starvation and heat stress. In males mutant for both *nrx-1* and *nlg-1*, mid-L4 starvation did not induce DVB neurite outgrowth compared with fed *nrx-1; nlg-1* double mutant males (**Fig.9A-C**), suggestive that NRX-1 and NLG-1 function together to promote adolescent starvation-induced DVB neurite outgrowth. This is in striking contrast to the role previously described for NLG-1 after day 1 of adulthood, where it antagonizes NRX-1 and represses DVB experience-dependent neurite outgrowth (Hart & Hobert, 2018). Perhaps at day 1 NRX-1 and NLG-1 provide adhesive connection required for early outgrowth and/or synapse stability, which later becomes antagonistic to balance outgrowth and synaptic strength. The shared requirement could also be from upstream of DVB at synapses required for neuronal signaling of starvation stress, or via a cell non-autonomous function, although previous examples of cell non-autonomous functions of NRX-1 were not dependent on NLG-1 (Philbrook et al., 2018).

**Figure 9.**
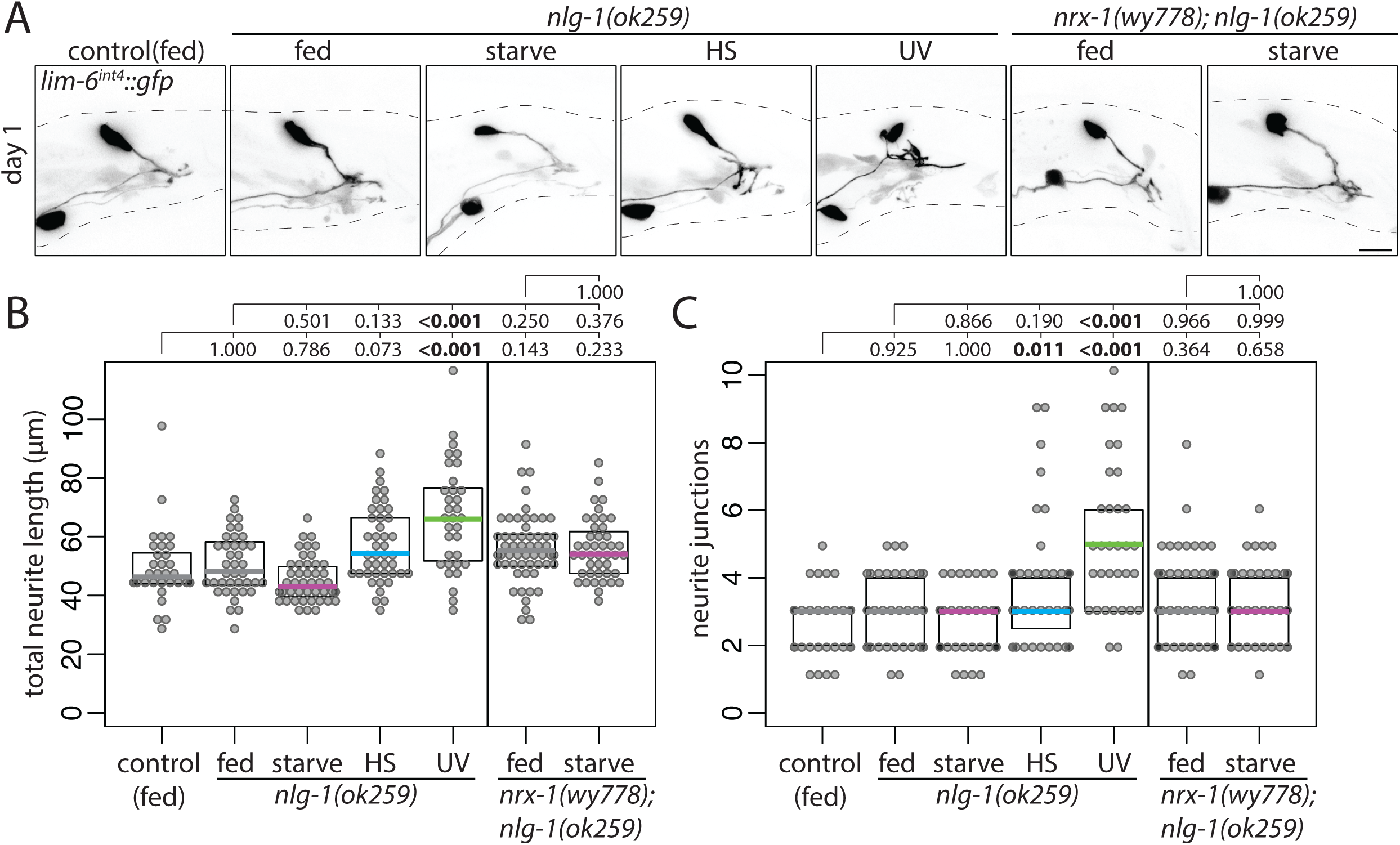
Neuroligin is required for DVB neurite outgrowth following adolescent starvation and heat stress. **A)** Confocal z-stack maximum projections of DVB neuron in fed control, fed and mid-L4 stressed *nlg-1(ok259)* males, and fed and mid-L4 starved *nrx-1(wy778)*; *nlg-1(ok259)* males at day 1 of adulthood (scale bar = 10μm). Quantification of (**B**) total neurite length and (**C**) neurite junctions for DVB in fed control, fed and mid-L4 stressed *nlg-1(ok259)* males, and fed and mid-L4 starved *nrx-1(wy778)*; *nlg-1(ok259)* males at day 1 of adulthood.

### Stress responsive transcription factors *daf-16* and *hsf-1* are required for starvation induced DVB neurite outgrowth

How is mid-L4 stress signaled to the DVB neuron to induce neurite outgrowth at day 1 of adulthood? In *C. elegans*, stress resistance, stress response, and the interplay between longevity and stress are signaled through two conserved transcription factors, *FOXO*/*daf-16* and the heat shock factor, *HSF1*/*hsf-1*, both with roles in conserved insulin signaling pathways (Baumeister, Schaffitzel, & Hertweck, 2006; Hsu, Murphy, & Kenyon, 2003). To test if these factors play a role in signaling stress to DVB, males mutant for *daf-16* or *hsf-1* were exposed to mid-L4 stress. UV exposure induced DVB neurite outgrowth in *daf-16* mutant males compared to unstressed *daf-16* males and control males. Heat stress did not induce neurite outgrowth in *daf-16* males, and starvation only partially induced neurite outgrowth in *daf-16* males (**Fig.10A-C**). In *hsf-1* mutant males, mid-L4 starvation did not increase DVB neurites compared with fed controls and unstressed *hsf-1* mutant males (**Fig.10D-F**). However, mid-L4 heat stress induced neurite outgrowth and UV stress partially induced neurite outgrowth in *hsf-1* mutant males (**Fig.10D-F**). Thus, *daf-16* and *hsf-1* are involved in DVB neurite outgrowth for some, but not all stressors, further highlighting the complexity of molecular mechanisms underlying neuronal remodeling in response to different stressors. The role of *daf-16* and *hsf-1* in DVB neurite outgrowth are unique to stress response, as neither, in the absence of stress, impacted experience-dependent DVB neurite outgrowth at day 3 (**Fig.11A-C**). *daf-16* and *hsf-1* have distinct and overlapping target genes, and each can regulate target genes in a tissue-specific manner that is not fully understood (Baumeister et al., 2006; Hsu et al., 2003; Volovik et al., 2014). Due to the complexity of these signaling systems in many tissues beyond DVB and neurons, further work will be needed to determine their precise location and tissues of action. One intriguing possibility is that *nrx-1/nlg-1* are targets of these factors in neurons, and there is some limited evidence that factors involved in insulin signaling pathways regulate *nrx-1* and *nlg-1* in certain circumstances (Shen, Wang, & Wang, 2007; Staab, Evgrafov, Knowles, & Sieburth, 2014).

**Figure 10.**
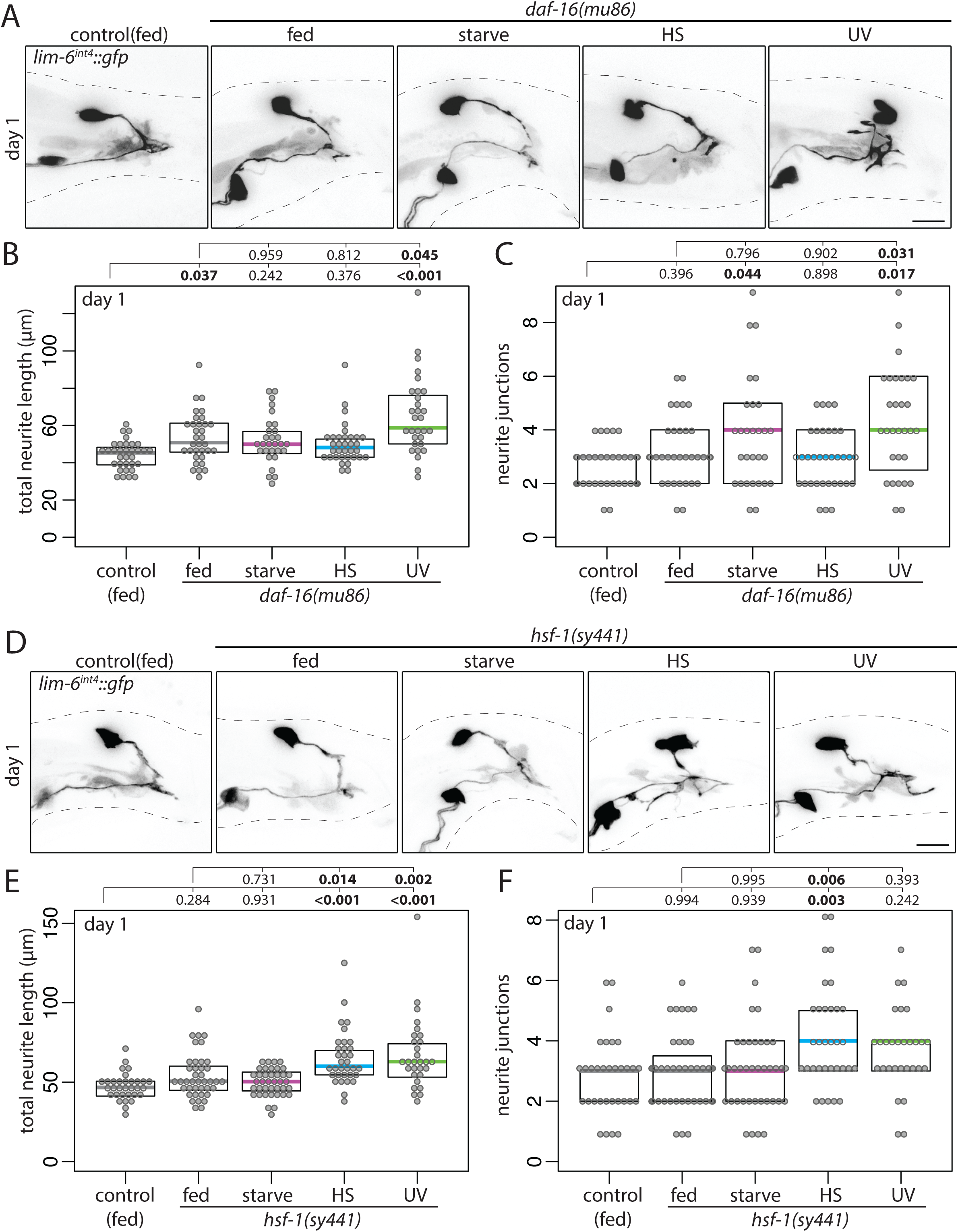
Stress responsive transcription factors DAF-16 and HSF-1 are required for DVB neurite outgrowth following adolescent starvation. **A)** Confocal z-stack maximum projections of DVB neuron in fed control and fed and mid-L4 stressed *daf-16(mu86)* males at day 1 of adulthood (scale bar = 10μm). Quantification of (**B**) total neurite length and (**C**) neurite junctions for DVB in fed control and fed and mid-L4 stressed *daf-16(mu86)* males at day 1. **D)** Confocal z-stack maximum projections of DVB neuron in fed control and fed and mid-L4 stressed *hsf-1(sy441)* males at day 1 of adulthood (scale bar = 10μm). Quantification of (**E**) total neurite length and (**F**) neurite junctions for DVB in fed and fed and mid-L4 stressed *hsf-1(sy441)* males at day 1.

**Figure 11.**
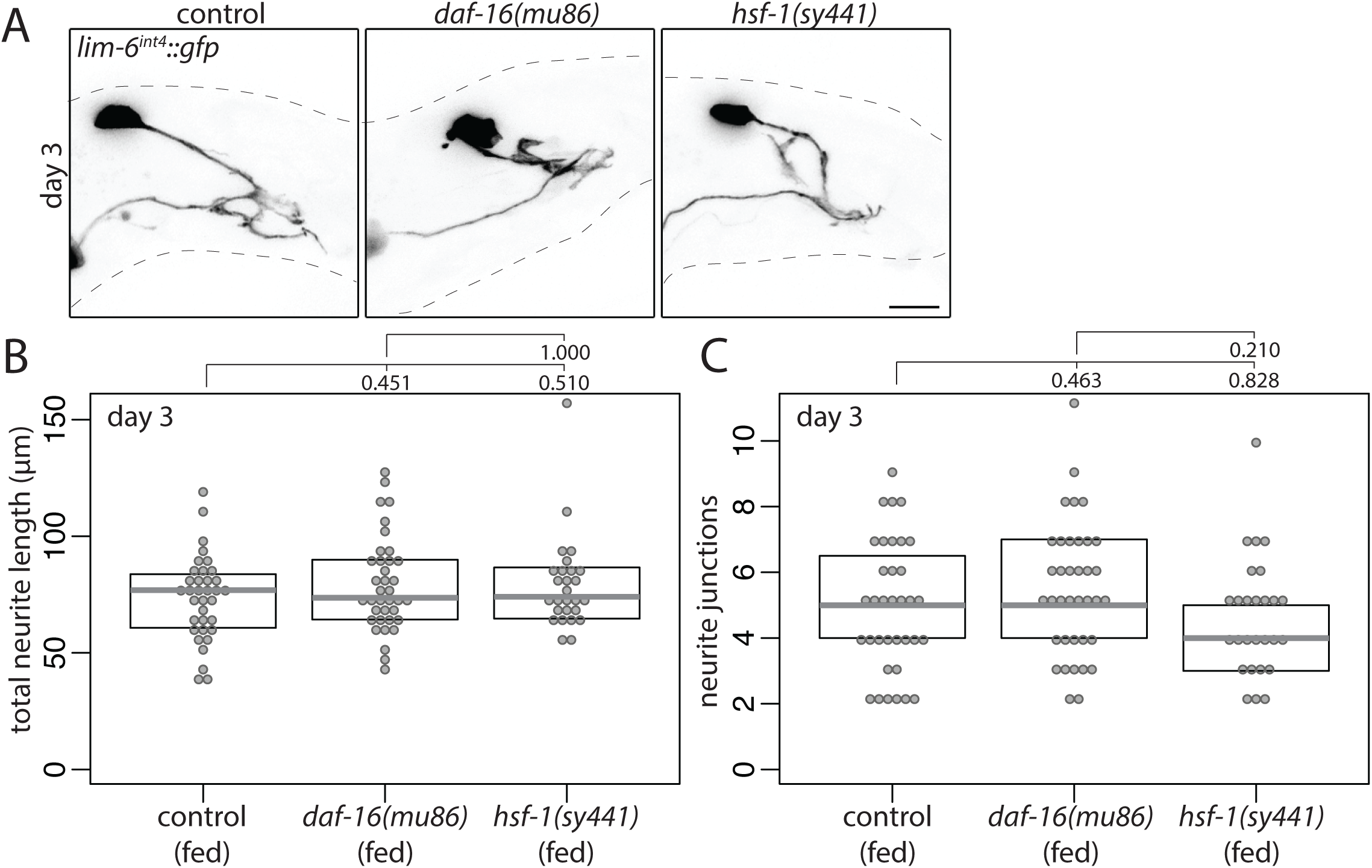
Stress responsive transcription factors DAF-16 and HSF-1 are not required for experience-dependent DVB neurite outgrowth in adulthood. **A)** Confocal z-stack maximum projections of DVB neuron in fed control, fed *daf-16(mu86)*, and fed *hsf-1(sy441)* males at day 3 of adulthood (scale bar = 10μm). Quantification of (**B**) total neurite length and (**C**) neurite junctions for DVB in fed control, fed *daf-16(mu86)*, and fed *hsf-1(sy441)* males at day 3.

## DISCUSSION

Neurons can respond to stress by altering the morphology of their dendritic trees, altering their connectivity with other neurons within circuits, and thus contributing to alterations in behavioral output (McEwen et al., 2016). Utilizing a recently-discovered form of experience-dependent morphologic plasticity in adult male *C. elegans*, I here report that stress induces morphologic remodeling of neurites of a GABAergic neuron and directly impacts behavior. In this study, I find that three stressors during adolescence result in altered morphologic plasticity in the form of increases in axon-like projections of a GABAergic neuron upon entry into adulthood. Remarkably, although each stressor induced increased neurite outgrowth, their effects on a specific behavior (i.e. spicule protraction) were different, which was further reflected in slight differences between stressors on long-term morphologic changes. Interestingly, it is only stressors with a functional impact on behavior that were found to be dependent on *nrx-1* (**Fig.12**); only nutritional stress resulted in both morphologic and functional changes that were dependent on neurexin. In contrast, genotoxic stress resulted in morphologic changes without any impact on behavior and was not dependent on *nrx-1* (**Fig.12**), while heat stress also resulted in morphologic changes not dependent on *nrx-1* and without any impact on behavior, but with loss of *nrx-1* uncovering a previously undetectable functional change in the circuit following heat stress. Further, when analyzing the role of neuroligin/*nlg-1* and stress responsive transcription factors *FOXO*/*daf-16* and *hsf-1*, it appears that in response to each stressor, a unique combination of molecular mechanisms control a similar pattern of remodeling (**Fig.12**). Together, these findings provide evidence that neuronal morphologic changes in response to stress are dynamic and that different types of stress can have different functional impacts and underlying molecular mechanisms, despite the observation that they all induce changes to neuron morphology. Furthermore, neuronal stress response at the functional level is more complex than a readout of morphologic changes, even in a simple nervous system.

**Figure 12.**
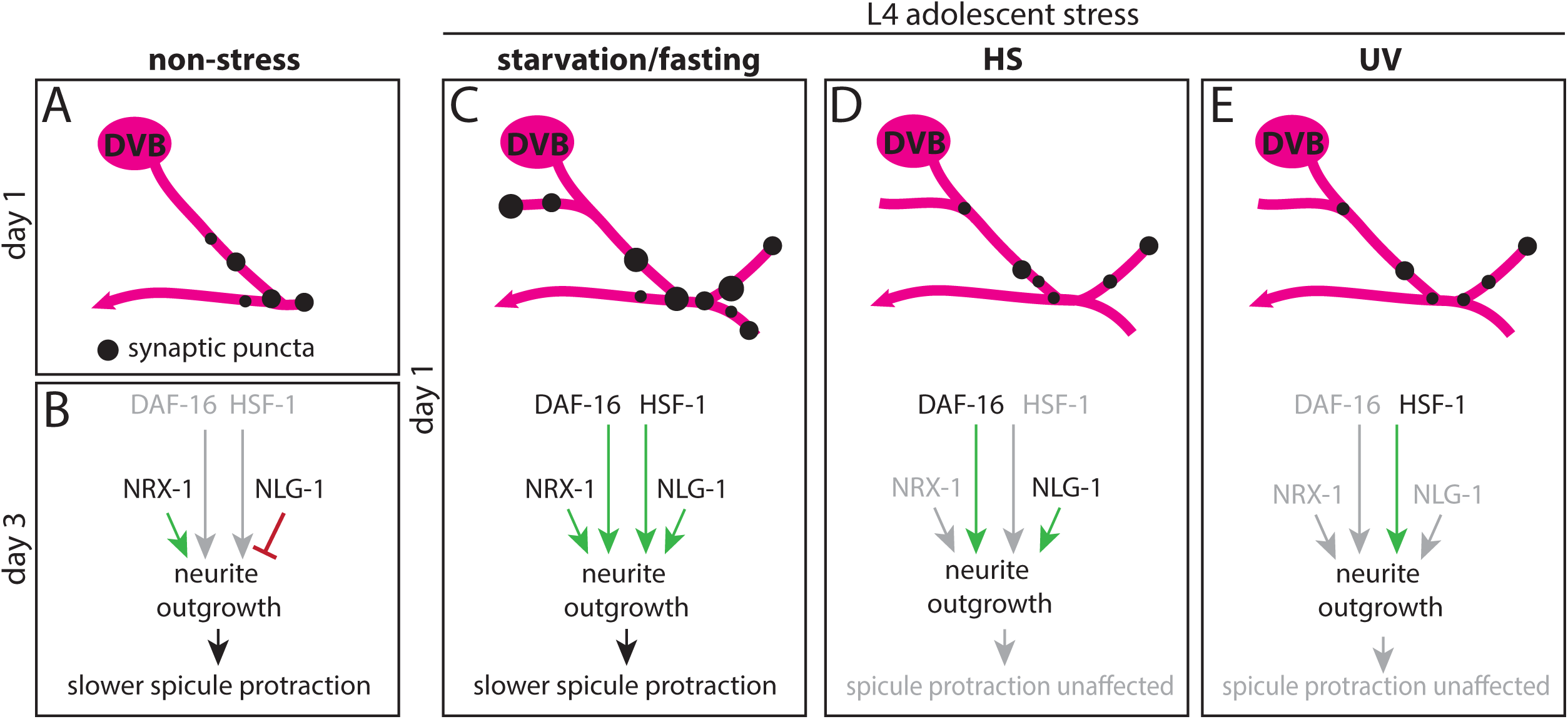
Model of DVB morphologic, synaptic, and behavioral response to adolescent stressors with differential underlying molecular mechanisms. **A)** DVB in non-stressed males at day 1 has few neurites and few presynaptic sites. **B)** After day 1, neurites will grow out in response to experience/activity, increasing number of presynaptic puncta and is dependent on NRX-1 in DVB, antagonized by NLG-1 in post-synaptic tissues, and results in slower spicule protraction at day 3. **C)** Adolescent starvation increases DVB neurite outgrowth at day 1 of adulthood with corresponding increases in pre-synaptic sites and slower spicule protraction. Starvation-induced neurites are dependent on NRX-1 and NLG-1, as well as the transcription factors DAF-16 and HSF-1. **D)** Adolescent exposure to heat stress increases DVB neurite outgrowth at day 1, dependent on NLG-1 and partly on DAF-16, but not NRX-1. This outgrowth does not increase pre-synaptic sites or alter spicule protraction behavior. **D)** Adolescent exposure to UV increases DVB neurite outgrowth at day 1 without increasing pre-synaptic sites or altering spicule protraction behavior. UV-induced DVB neurite outgrowth is not dependent on NRX-1, NLG-1, but is partly dependent on HSF-1.

It was recently reported that past life experience, specifically starvation stress, impacts the development of male-specific nervous system wiring in *C. elegans* through monoamine signaling (Bayer & Hobert, 2018). Starvation during development before (L1-L3), but not during adolescence (L4), blocks synaptic pruning events that are required for male-specific circuit wiring and behavior (Bayer & Hobert, 2018; Oren-Suissa, Bayer, & Hobert, 2016). Pre-adolescent starvation stress produced a ‘memory’ of starvation in the form of sex-specific connections not being pruned. Interestingly, the results reported here demonstrate that starvation stress during adolescence produces changes in male-specific connectivity through a different mechanism, as a result of altering GABAergic morphologic plasticity. In combination, these results show that early life and adolescent stress both influence the connectivity and behavioral output of sexually dimorphic circuits, the former by impacting the development of sex-specific neuronal connectivity, and the latter by altering the dynamics of an adult form of experience-dependent morphologic plasticity.

Early-life and adolescence are important periods in brain development and maturation marked by reorganization of neurons and circuits in the brain, thereby making these periods sensitive to stress disruption. In adolescence, stress has been shown to impact dendritic morphology of neurons in a number of brain regions, in some cases having a unique impact on morphology and behavior in adolescence compared to the same stress during adulthood (Tsai et al., 2014; Tzanoulinou et al., 2014), with stresses during critical periods in brain development linked to neuropsychiatric disorders. A two-hit model has been proposed where stress inflicted on a genetically-predisposed individual during critical brain development windows (early brain development and adolescent maturation) plays a role in the emergence of symptoms. Interestingly, there is evidence that neuron morphology and relevant behavioral phenotypes are impacted differently in animals stressed during both early life and adolescent critical periods of brain development compared to stress during one critical period or in adulthood (Eiland & McEwen, 2012). The results reported here demonstrate that even in the nematode *C. elegans*, adolescence and sexual maturation of the nervous system are a period of particular sensitivity to stress.

It has been proposed that to develop models of severe psychiatric disorders, studies should combine genetic variants in susceptibility genes with several stressors during critical periods (Schmitt et al., 2014). Recent work has identified many genetic risk factors for ASDs and schizophrenia that converge on shared pathways, some of which could potentially enhance sensitivity of the nervous system to stress. Neurexin and neuroligin have been associated genetically with ASDs and schizophrenia, but their relationships to neuronal stress response have not been studied. The results reported here indicate that neurexin/neuroligin play a role in neuronal stress response and promotes morphologic remodeling of GABAergic axons in response to stress. Another adhesion molecule, *NCAM*, has a described role in stress-induced neuronal remodeling, suggesting that multiple cellular adhesion molecules may be important in this process (Gilabert-Juan et al., 2011; McCall et al., 2013). A very recent report finds another ASD-associated gene, *ANK2*, regulates neuronal branching patterns in mice, through an interaction with an axonal adhesion molecule, *L1CAM*, further implicating neuron morphology, resulting connectivity defects, and adhesion molecules in pathogenesis of neurodevelopmental disorders (Yang et al., 2019). The pathogenic role of *NRXN1* in ASDs and schizophrenia has been complicated by the presence of deletions/mutations in unaffected parents and siblings, suggesting variable penetrance or involvement of *NRXN1* in these disorders (Lowther et al., 2017; Todarello et al., 2014; Woodbury-Smith et al., 2017). The results reported here in *C. elegans* describe the impact of alteration in a single gene on the morphologic and behavioral consequences of different stress exposures on a single neuron, and suggest that some phenotypic heterogeneity may potentially be explained by the requirement for a genetic risk to be uncovered by a specifically timed stress. In terms of neuropsychiatric disease, perhaps multiple genetic risk factors, in combination with different environmental exposures act together to impact the complex human brain, but with varying phenotypic consequences depending on the specific genetic/environmental combination. In fact, a recent study suggests that variation in genetic background in combination with defects in developmental susceptibility genes could explain phenotypic variability (Pizzo et al., 2018), and perhaps stress could also be added to this model. Underlying genetic differences, including defects in susceptibility genes, may only impact some neuronal responses to stress during certain critical periods, which may differ for each stress/genetic risk, adding further complexity and variability to relevant phenotypes.

## ACKNOWLEDGEMENTS

I thank Meera Sundaram, David Raizen, Oliver Hobert, Emily Bayer, Laura Pereira, and Theodore G. Drivas for discussions and comments on the included experiments and manuscript; and Meera Sundaram and members of the Sundaram lab for technical support. M.P.H. is supported in part by the Autism Spectrum Program of Excellence at the Perelman School of Medicine. Some strains were provided by the CGC, funded by NIH Office of Research Infrastructure Programs (P40 OD010440).

## Author Contributions

M.P.H. conceived, designed, and performed the experiments; analyzed data, and wrote the manuscript.

## Author Information

The author declares no conflicts of interest.

## MATERIALS AND METHODS

### *C. elegans* strains

Wild-type strains were *C. elegans* variety Bristol, strain N2. Worms were grown at 23 °C on nematode growth media (NGM) plates seeded with bacteria (*E.coli* OP50) as a food source. All males contained either *him-8(e1489) IV* or *him-5(e1490) V* as indicated by strain.

Mutant alleles used in this study include:

*him-8(e1489) IV, him-5(e1490) V, unc-119(ed3) III; nrx-1(wy778*[*unc-119(+)*]*) V, nrx-1(gk246237) V, nlg-1*(ok259) *X*, *daf-16(mu86) I, hsf-1(sy441) I*.

All transgenic strains used in this study are listed in Table 1 ordered by Figures. All plasmids were injected at 25 ng µl^−1^ with coinjection marker *myo-3::gfp* at 25 ng µl^−1^ to generate extrachromosomal arrays.

**Table 1.** Strain table

| Fig | Strain | mutant background | Array | DNA on array |
| --- | --- | --- | --- | --- |
| 1 | OH15098 | *him-8(e1489)* | *otIs525* | *lim-6<sup>int4</sup>::gfp* |
| 4 | MPH1 | *him-5(e1490)* | *otIs541* | *lim-6<sup>int4</sup>::wCherry* |
|  |  |  | *hpmEx1* | *lim-6<sup>int4</sup>::cla-1::gfp; myo-3::gfp* |
| 5 | OH15099 | *him-5(e1490)* | *otIs541* | see above |
|  |  |  | *otIs659* | *lim-6<sup>int4</sup>::gfp::rab-3; ttx-3::gfp* |
| 6 | OH15111 | *unc-119(ed3); nrx-1(wy778[unc-119(+)]); him-8(e1489)* | *otIs525* | see above |
| 6 | OH15116 | *nrx-1(gk246237); him-8(e1489)* | *otIs525* | see above |
| 8 | OH15112 | *unc-119(ed3); nrx-1(wy778[unc-119(+)]); him-8(e1489)* | *otIs525* | see above |
|  |  |  | *otEx7013* | *lim-6<sup>int4</sup>::birA::nrx-1* |
| 8 | MPH2 | *unc-119(ed3); nrx-1(wy778[unc-119(+)]); him-8(e1489)* | *otIs525* | see above |
|  |  |  | *hpmEx2* | *ric-19<sup>prom6</sup>::birA::nrx-1* |
| 9 | MPH3 | *nlg-1(ok259); him-8(e1489)* | *otIs525* | see above |
| 9 | OH15114 | *unc-119(ed3); nrx-1(wy778[unc-119(+)]); nlg-1(ok259); him-8(e1489)* | *otIs525* | see above |
| 10 | MPH4 | *daf-16(mu86); him-8(e1489)* | *otIs525* | see above |
| 10 | MPH5 | *hsf-1(sy441); him-8(e1489)* | *otIs525* | see above |

### Cloning and constructs

To generate *lim-6^int4^::cla-1::gfp* (pMPH21) a 291 bp fragment of the *lim-6* fourth intron was amplified with primers fwd - GATTACGCCAAGCTTGCATGCGGATCCTTAGCCAGTTGCATAA and rev – GAGGCGCGCCAATCCCGGGGATCCCCTGAAAATGTTCTATG and cloned into PK068 (a gift from Peri Kurshan (Kurshan et al., 2018)) to replace the *unc-17* promoter using RF cloning.

To generate *ric-19^prom6^::BirA::nrx-1 (*pMPH24), a 147 bp fragment of the *ric-19* promoter was PCR amplified from genomic DNA using primers fwd - GAAATGAAATAAAGCTTGCATGCATTAAAGAGTGTGCTCCAC and rev - CTTTGGGTCCTTTGGCCAATCCCGGGTTCAAAGTGAAGAGCTCTCTCGAC, and cloned into pMH27 (*lim-6^int4^::BirA::nrx-1)* to replace the *lim-6^int4^* promoter using RF cloning.

### Stress induction

Male worms were picked at early-L4, mid-L4, or following L4 molt (adulthood) as indicated in Figures onto plates described below. Non-stressed males were placed onto NGM plates with OP50 as a food source at 23°C. Starved and fasted males were placed onto NGM plates lacking peptone and any food source at times indicated and for duration indicated (18 hours or 4 hours), then analyzed or returned to NGM plates with OP50 as a food source. Heat stress was performed by placing males onto NGM plates with OP50 as a food source, setting the plate in a 37°C incubator for 30 minutes, then returning plate to 23°C. For ultraviolet (UV) exposure males were placed on NGM plates with a thin layer of OP50 as a food source to minimize blocking UV rays, which was set in a Spectrolinker XL-1500 (Spectroline) uncovered, and irradiated with 254 nm light using the energy input function which varies the exposure time (DeBardeleben, Lopes, Nessel, & Raizen, 2017), here set to 200 X100 μJ/cm^2^. In all non-stress and stress conditions, males were subjected to image analysis or behavioral assays at times indicated in Figures.

### Microscopy

Worms were anesthetized using 5 μl of 100 mM of sodium azide (NaN3) on a pad of 5% agarose on glass slides, and covered with a glass coverslip. Worms were analyzed by fluorescence microscopy, using an upright stand Leica TCS SP8 laser-scanning microscope operated by LAS X software. Confocal laser and photomultiplier tube settings were set at identical levels for image acquisition of all experimental conditions. Confocal z-stacks were reconstructed as maximum intensity projections in FIJI. Figures were prepared using Adobe Photoshop CS6 and Adobe Illustrator CS6.

### Neurite tracing

Neurite tracing was performed as previously described (Hart & Hobert, 2018). Briefly, confocal z-stacks were opened in FIJI, and analyzed using the Simple Neurite Tracer plugin (Longair, Baker, & Armstrong, 2011). The primary neurite of DVB was traced from the center of the cell soma to where the axon projects ventrally and turns anteriorly into the ventral nerve cord. Neurite branches were added by tracing off the primary neurite, including all neurites emanating posterior of the last branch point. The simple neurite tracer plugin was used to analyze the skeletons for neurite length, which were summed to calculate total neurite length, and the number of neurite junctions (a proxy for the number of neurite branches).

### Aldicarb spicule protraction assay

Aldicarb solution was added on the top of NGM agar plates and allowed to dry for 1 hour with plate lid on. For a concentration of 0.8 mM aldicarb, 80 ul of a 100 mM in 70% ethanol stock solution was added onto 10 ml NGM plates and spread evenly on surface (Locke et al., 2008). 10 male worms were placed onto the aldicarb plates and observed for spicule protraction longer than 3s, when the time was recorded for each worm. Aldicarb assay was performed on 2 technical replicates and at least 3 independent replicates. Investigator was blinded to experimental conditions before aldicarb assay was performed.

### Quantification of CLA-1::GFP puncta and particle analysis of GFP::RAB-3 synaptic puncta

Confocal z-stacks were opened in FIJI and CLA-1::GFP puncta overlapping with DVB neurites (labeled with mCherry) were counted by scanning through the stack for distinct puncta. Counting puncta were included on any neurites, minus the DVB cell body, up to the last branch point before the DVB projection turns anteriorly along ventral nerve cord, the same as the cutoff for neurite tracing. Confocal z-stacks of GFP::RAB-3 micrographs were opened and turned into maximum projections in FIJI. A region of interest was drawn around all of the neurites of DVB, defined the same way as for neurite tracing. Particles, representing synaptic puncta, were defined by manual thresholding of projections (top slider 20, bottom slider 255) and subsequent particle analysis to outline particles. Without applying threshold, these regions were then analyzed using Analyze Particle tool. Particle number, particle area, and mean particle intensity were determined for each particle and used to determine total particle number, mean particle area, and mean particle intensity for each worm.

### Statistics and reproducibility

One-way ANOVA with post-hoc Tukey HSD tests were performed using *RStudio*, p-values shown on each graph. Beeswarm and boxplot graphs were created in *RStudio*.

## REFERENCES

Autism Spectrum Disorders Working Group of The Psychiatric Genomics, C. (2017). Meta-analysis of GWAS of over 16,000 individuals with autism spectrum disorder highlights a novel locus at 10q24.32 and a significant overlap with schizophrenia. Mol Autism, 8, 21. doi:10.1186/s13229-017-0137-9

Baumeister, R., Schaffitzel, E., & Hertweck, M. (2006). Endocrine signaling in Caenorhabditis elegans controls stress response and longevity. J Endocrinol, 190(2), 191–202. doi:10.1677/joe.1.06856

Bayer, E. A., & Hobert, O. (2018). Past experience shapes sexually dimorphic neuronal wiring through monoaminergic signalling. Nature, 561(7721), 117–121. doi:10.1038/s41586-018-0452-0

Bishop-Fitzpatrick, L., Mazefsky, C. A., Minshew, N. J., & Eack, S. M. (2015). The relationship between stress and social functioning in adults with autism spectrum disorder and without intellectual disability. Autism Res, 8(2), 164–173. doi:10.1002/aur.1433

Cattane, N., Richetto, J., & Cattaneo, A. (2018). Prenatal exposure to environmental insults and enhanced risk of developing Schizophrenia and Autism Spectrum Disorder: focus on biological pathways and epigenetic mechanisms. Neurosci Biobehav Rev. doi:10.1016/j.neubiorev.2018.07.001

Chaste, P., & Leboyer, M. (2012). Autism risk factors: genes, environment, and gene-environment interactions. Dialogues Clin Neurosci, 14(3), 281–292.

Conrad, C. D. (2006). What is the functional significance of chronic stress-induced CA3 dendritic retraction within the hippocampus? Behav Cogn Neurosci Rev, 5(1), 41–60. doi:10.1177/1534582306289043

Conrad, C. D., LeDoux, J. E., Magarinos, A. M., & McEwen, B. S. (1999). Repeated restraint stress facilitates fear conditioning independently of causing hippocampal CA3 dendritic atrophy. Behav Neurosci, 113(5), 902–913.

Conrad, C. D., Ortiz, J. B., & Judd, J. M. (2017). Chronic stress and hippocampal dendritic complexity: Methodological and functional considerations. Physiol Behav, 178, 66–81. doi:10.1016/j.physbeh.2016.11.017

Cook, S. C., & Wellman, C. L. (2004). Chronic stress alters dendritic morphology in rat medial prefrontal cortex. J Neurobiol, 60(2), 236–248. doi:10.1002/neu.20025

Czeh, B., Vardya, I., Varga, Z., Febbraro, F., Csabai, D., Martis, L. S., … Wiborg, O. (2018). Long-Term Stress Disrupts the Structural and Functional Integrity of GABAergic Neuronal Networks in the Medial Prefrontal Cortex of Rats. Front Cell Neurosci, 12, 148. doi:10.3389/fncel.2018.00148

DeBardeleben, H. K., Lopes, L. E., Nessel, M. P., & Raizen, D. M. (2017). Stress-Induced Sleep After Exposure to Ultraviolet Light Is Promoted by p53 in Caenorhabditis elegans. Genetics, 207(2), 571–582. doi:10.1534/genetics.117.300070

Eiland, L., & McEwen, B. S. (2012). Early life stress followed by subsequent adult chronic stress potentiates anxiety and blunts hippocampal structural remodeling. Hippocampus, 22(1), 82–91. doi:10.1002/hipo.20862

Fuld, S. (2018). Autism Spectrum Disorder: The Impact of Stressful and Traumatic Life Events and Implications for Clinical Practice. Clin Soc Work J, 46(3), 210–219. doi:10.1007/s10615-018-0649-6

Garcia, L. R., Mehta, P., & Sternberg, P. W. (2001). Regulation of distinct muscle behaviors controls the C. elegans male’s copulatory spicules during mating. Cell, 107(6), 777–788.

Gauthier, J., Siddiqui, T. J., Huashan, P., Yokomaku, D., Hamdan, F. F., Champagne, N., … Rouleau, G. A. (2011). Truncating mutations in NRXN2 and NRXN1 in autism spectrum disorders and schizophrenia. Hum Genet, 130(4), 563–573. doi:10.1007/s00439-011-0975-z

Gilabert-Juan, J., Bueno-Fernandez, C., Castillo-Gomez, E., & Nacher, J. (2017). Reduced interneuronal dendritic arborization in CA1 but not in CA3 region of mice subjected to chronic mild stress. Brain Behav, 7(2), e00534. doi:10.1002/brb3.534

Gilabert-Juan, J., Castillo-Gomez, E., Guirado, R., Molto, M. D., & Nacher, J. (2013). Chronic stress alters inhibitory networks in the medial prefrontal cortex of adult mice. Brain Struct Funct, 218(6), 1591–1605. doi:10.1007/s00429-012-0479-1

Gilabert-Juan, J., Castillo-Gomez, E., Perez-Rando, M., Molto, M. D., & Nacher, J. (2011). Chronic stress induces changes in the structure of interneurons and in the expression of molecules related to neuronal structural plasticity and inhibitory neurotransmission in the amygdala of adult mice. Exp Neurol, 232(1), 33–40. doi:10.1016/j.expneurol.2011.07.009

Guo, X., Navetta, A., Gualberto, D. G., & Garcia, L. R. (2012). Behavioral decay in aging male C. elegans correlates with increased cell excitability. Neurobiol Aging, 33(7), 1483 e1485-1423. doi:10.1016/j.neurobiolaging.2011.12.016

Hallmayer, J., Cleveland, S., Torres, A., Phillips, J., Cohen, B., Torigoe, T., … Risch, N. (2011). Genetic heritability and shared environmental factors among twin pairs with autism. Arch Gen Psychiatry, 68(11), 1095–1102. doi:10.1001/archgenpsychiatry.2011.76

Hart, M. P., & Hobert, O. (2018). Neurexin controls plasticity of a mature, sexually dimorphic neuron. Nature, 553(7687), 165–170. doi:10.1038/nature25192

Hertz-Picciotto, I., Schmidt, R. J., & Krakowiak, P. (2018). Understanding environmental contributions to autism: Causal concepts and the state of science. Autism Res, 11(4), 554–586. doi:10.1002/aur.1938

Hsu, A. L., Murphy, C. T., & Kenyon, C. (2003). Regulation of aging and age-related disease by DAF-16 and heat-shock factor. Science, 300(5622), 1142–1145. doi:10.1126/science.1083701

Kagias, K., Nehammer, C., & Pocock, R. (2012). Neuronal responses to physiological stress. Front Genet, 3, 222. doi:10.3389/fgene.2012.00222

Kirov, G., Rujescu, D., Ingason, A., Collier, D. A., O’Donovan, M. C., & Owen, M. J. (2009). Neurexin 1 (NRXN1) deletions in schizophrenia. Schizophr Bull, 35(5), 851–854. doi:10.1093/schbul/sbp079

Kurshan, P. T., Merrill, S. A., Dong, Y., Ding, C., Hammarlund, M., Bai, J., … Shen, K. (2018). gamma-Neurexin and Frizzled Mediate Parallel Synapse Assembly Pathways Antagonized by Receptor Endocytosis. Neuron, 100(1), 150–166 e154. doi:10.1016/j.neuron.2018.09.007

LeBoeuf, B., Guo, X., & Garcia, L. R. (2011). The effects of transient starvation persist through direct interactions between CaMKII and ether-a-go-go K+ channels in C. elegans males. Neuroscience, 175, 1–17. doi:10.1016/j.neuroscience.2010.12.002

Locke, C., Berry, K., Kautu, B., Lee, K., Caldwell, K., & Caldwell, G. (2008). Paradigms for pharmacological characterization of C. elegans synaptic transmission mutants. J Vis Exp(18). doi:10.3791/837

Longair, M. H., Baker, D. A., & Armstrong, J. D. (2011). Simple Neurite Tracer: open source software for reconstruction, visualization and analysis of neuronal processes. Bioinformatics, 27(17), 2453–2454. doi:10.1093/bioinformatics/btr390

Lowther, C., Speevak, M., Armour, C. M., Goh, E. S., Graham, G. E., Li, C., … Bassett, A. S. (2017). Molecular characterization of NRXN1 deletions from 19,263 clinical microarray cases identifies exons important for neurodevelopmental disease expression. Genet Med, 19(1), 53–61. doi:10.1038/gim.2016.54

Maro, G. S., Gao, S., Olechwier, A. M., Hung, W. L., Liu, M., Ozkan, E., … Shen, K. (2015). MADD-4/Punctin and Neurexin Organize C. elegans GABAergic Postsynapses through Neuroligin. Neuron, 86(6), 1420–1432. doi:10.1016/j.neuron.2015.05.015

McCall, T., Weil, Z. M., Nacher, J., Bloss, E. B., El Maarouf, A., Rutishauser, U., & McEwen, B. S. (2013). Depletion of polysialic acid from neural cell adhesion molecule (PSA-NCAM) increases CA3 dendritic arborization and increases vulnerability to excitotoxicity. Exp Neurol, 241, 5–12. doi:10.1016/j.expneurol.2012.11.028

McEwen, B. S., Nasca, C., & Gray, J. D. (2016). Stress Effects on Neuronal Structure: Hippocampus, Amygdala, and Prefrontal Cortex. Neuropsychopharmacology, 41(1), 3–23. doi:10.1038/npp.2015.171

McKlveen, J. M., Morano, R. L., Fitzgerald, M., Zoubovsky, S., Cassella, S. N., Scheimann, J. R., … Herman, J. P. (2016). Chronic Stress Increases Prefrontal Inhibition: A Mechanism for Stress-Induced Prefrontal Dysfunction. Biol Psychiatry, 80(10), 754–764. doi:10.1016/j.biopsych.2016.03.2101

Oren-Suissa, M., Bayer, E. A., & Hobert, O. (2016). Sex-specific pruning of neuronal synapses in Caenorhabditis elegans. Nature, 533(7602), 206–211. doi:10.1038/nature17977

Philbrook, A., Ramachandran, S., Lambert, C. M., Oliver, D., Florman, J., Alkema, M. J., … Francis, M. M. (2018). Neurexin directs partner-specific synaptic connectivity in C. elegans. Elife, 7. doi:10.7554/eLife.35692

Pizzo, L., Jensen, M., Polyak, A., Rosenfeld, J. A., Mannik, K., Krishnan, A., … Girirajan, S. (2018). Rare variants in the genetic background modulate cognitive and developmental phenotypes in individuals carrying disease-associated variants. Genet Med. doi:10.1038/s41436-018-0266-3

Reichelt, A. C., Rodgers, R. J., & Clapcote, S. J. (2012). The role of neurexins in schizophrenia and autistic spectrum disorder. Neuropharmacology, 62(3), 1519–1526. doi:10.1016/j.neuropharm.2011.01.024

Sandin, S., Lichtenstein, P., Kuja-Halkola, R., Larsson, H., Hultman, C. M., & Reichenberg, A. (2014). The familial risk of autism. JAMA, 311(17), 1770–1777. doi:10.1001/jama.2014.4144

Schmitt, A., Malchow, B., Hasan, A., & Falkai, P. (2014). The impact of environmental factors in severe psychiatric disorders. Front Neurosci, 8, 19. doi:10.3389/fnins.2014.00019

Shen, L. L., Wang, Y., & Wang, D. Y. (2007). Involvement of genes required for synaptic function in aging control in C. elegans. Neurosci Bull, 23(1), 21–29. doi:10.1007/s12264-007-0003-4

Snoek, L. B., Sterken, M. G., Volkers, R. J., Klatter, M., Bosman, K. J., Bevers, R. P., … Kammenga, J. E. (2014). A rapid and massive gene expression shift marking adolescent transition in C. elegans. Sci Rep, 4, 3912. doi:10.1038/srep03912

Staab, T. A., Evgrafov, O., Knowles, J. A., & Sieburth, D. (2014). Regulation of synaptic nlg-1/neuroligin abundance by the skn-1/Nrf stress response pathway protects against oxidative stress. PLoS Genet, 10(1), e1004100. doi:10.1371/journal.pgen.1004100

Stefanakis, N., Carrera, I., & Hobert, O. (2015). Regulatory Logic of Pan-Neuronal Gene Expression in C. elegans. Neuron, 87(4), 733–750. doi:10.1016/j.neuron.2015.07.031

Sudhof, T. C. (2017). Synaptic Neurexin Complexes: A Molecular Code for the Logic of Neural Circuits. Cell, 171(4), 745–769. doi:10.1016/j.cell.2017.10.024

Tick, B., Bolton, P., Happe, F., Rutter, M., & Rijsdijk, F. (2016). Heritability of autism spectrum disorders: a meta-analysis of twin studies. J Child Psychol Psychiatry, 57(5), 585–595. doi:10.1111/jcpp.12499

Todarello, G., Feng, N., Kolachana, B. S., Li, C., Vakkalanka, R., Bertolino, A., … Straub, R. E. (2014). Incomplete penetrance of NRXN1 deletions in families with schizophrenia. Schizophr Res, 155(1-3), 1–7. doi:10.1016/j.schres.2014.02.023

Tong, X. J., Hu, Z., Liu, Y., Anderson, D., & Kaplan, J. M. (2015). A network of autism linked genes stabilizes two pools of synaptic GABA(A) receptors. Elife, 4, e09648. doi:10.7554/eLife.09648

Tong, X. J., Lopez-Soto, E. J., Li, L., Liu, H., Nedelcu, D., Lipscombe, D., … Kaplan, J. M. (2017). Retrograde Synaptic Inhibition Is Mediated by alpha-Neurexin Binding to the alpha2delta Subunits of N-Type Calcium Channels. Neuron, 95(2), 326–340 e325. doi:10.1016/j.neuron.2017.06.018

Tsai, S. F., Huang, T. Y., Chang, C. Y., Hsu, Y. C., Chen, S. J., Yu, L., … Jen, C. J. (2014). Social instability stress differentially affects amygdalar neuron adaptations and memory performance in adolescent and adult rats. Front Behav Neurosci, 8, 27. doi:10.3389/fnbeh.2014.00027

Tu, H., Pinan-Lucarre, B., Ji, T., Jospin, M., & Bessereau, J. L. (2015). C. elegans Punctin Clusters GABA(A) Receptors via Neuroligin Binding and UNC-40/DCC Recruitment. Neuron, 86(6), 1407–1419. doi:10.1016/j.neuron.2015.05.013

Tzanoulinou, S., Garcia-Mompo, C., Castillo-Gomez, E., Veenit, V., Nacher, J., & Sandi, C. (2014). Long-term behavioral programming induced by peripuberty stress in rats is accompanied by GABAergic-related alterations in the Amygdala. PLoS One, 9(4), e94666. doi:10.1371/journal.pone.0094666

Volovik, Y., Moll, L., Marques, F. C., Maman, M., Bejerano-Sagie, M., & Cohen, E. (2014). Differential regulation of the heat shock factor 1 and DAF-16 by neuronal nhl-1 in the nematode C. elegans. Cell Rep, 9(6), 2192–2205. doi:10.1016/j.celrep.2014.11.028

Vyas, A., Mitra, R., Shankaranarayana Rao, B. S., & Chattarji, S. (2002). Chronic stress induces contrasting patterns of dendritic remodeling in hippocampal and amygdaloid neurons. J Neurosci, 22(15), 6810–6818. doi:20026655

Waltereit, R., Banaschewski, T., Meyer-Lindenberg, A., & Poustka, L. (2014). Interaction of neurodevelopmental pathways and synaptic plasticity in mental retardation, autism spectrum disorder and schizophrenia: implications for psychiatry. World J Biol Psychiatry, 15(7), 507–516. doi:10.3109/15622975.2013.838641

Woodbury-Smith, M., Nicolson, R., Zarrei, M., Yuen, R. K. C., Walker, S., Howe, J., … Scherer, S. W. (2017). Variable phenotype expression in a family segregating microdeletions of the NRXN1 and MBD5 autism spectrum disorder susceptibility genes. NPJ Genom Med, 2. doi:10.1038/s41525-017-0020-9

Xuan, Z., Manning, L., Nelson, J., Richmond, J. E., Colon-Ramos, D. A., Shen, K., & Kurshan, P. T. (2017). Clarinet (CLA-1), a novel active zone protein required for synaptic vesicle clustering and release. Elife, 6. doi:10.7554/eLife.29276

Yang, R., Walder-Christensen, K. K., Kim, N., Wu, D., Lorenzo, D. N., Badea, A., … Bennett, V. (2019). ANK2 autism mutation targeting giant ankyrin-B promotes axon branching and ectopic connectivity. Proc Natl Acad Sci U S A, 116(30), 15262–15271. doi:10.1073/pnas.1904348116

